# ResXR: infrastructure for quality-controlled, reproducible studies of human behavior in Extended Reality

**DOI:** 10.64898/2026.06.17.732869

**Authors:** Yehuda Bergstein, Noa Barel, Galia Shai Basson, Omri Bromberg, Tom Schonberg

## Abstract

Extended Reality (XR) combines experimental control and ecological validity, yet behavioral XR research lacks shared infrastructure: building immersive experiments demands specialized engineering, and custom tools yield data in custom formats that other laboratories cannot readily reanalyze. We present ResXR (Research with XR), an open-source toolkit providing a path from immersive experiment to standardized dataset and quality report, running on standalone Meta Quest headsets. A Unity template records head, hand, eye, and face tracking with per-sample software development kit (SDK) timestamps and status flags; an independent Python pipeline checks data quality, can mask flagged intervals, and exports raw and derivative datasets in Motion-BIDS, the motion extension of the Brain Imaging Data Structure (BIDS), with self-contained quality reports. Three ready-to-run paradigms, covering discrete choice, spatial navigation, and free visual exploration, let researchers collect and process data without programming. ResXR grounds its default schema and the signals its quality checks read in stress tests rather than in vendor documentation: when we induced tracking failures on Meta Quest Pro headsets, several status flags kept marking the data as valid. With its code, stress-test protocols and example datasets openly available, ResXR provides a transparent, community-extensible foundation for reproducible XR experimentation.

## Introduction

Over the past 30 years, extended reality (XR), encompassing virtual reality (VR), augmented reality (AR), and mixed reality (MR), has transitioned from specialized laboratories to consumer platforms, catalyzing substantial growth in experimental research. Analysis of over 21,000 experimental XR publications from 1992 to mid-2024 shows that more than half appeared in the last six years of that period, with growth accelerating after consumer headsets such as the Oculus Rift were released in the mid-2010s (Bailenson et al., 2025; Markowitz & Bailenson, 2025). XR’s appeal stems from its unique combination of experimental control and ecological validity: as an immersive system it elicits physiological, emotional, and behavioral responses that differ from two-dimensional paradigms, letting researchers observe realistic behavior within controlled digital environments (Hepperle & Wölfel, 2023; Pan & Hamilton, 2018; Wrzus et al., 2024). Around 2020, according to a timeline in Bailenson et al. (2025), barriers to entry lessened as researchers converged on Meta headsets and Unity software. Yet this growth has not translated into mainstream psychological research. The share of psychology articles in PubMed that include immersive VR declined from 0.2% to 0.05% between 2013 and 2023, and basic research on cognitive, emotional, and behavioral processes remains particularly scarce (Wrzus et al., 2024). A shared headset and engine lowered the cost of entry, but not the cost of building a sound behavioral measurement.

We identified three main barriers. First, in sensor-based measurement, invalid data can look valid: unlike missing data, they are not necessarily marked in the recording and have to be identified through quality checks (e.g., Mathôt & Vilotijević, 2023, for pupillometry). A device can continue to record after it has stopped measuring the participant, or record a signal that did not originate in the process of interest, and return values that resemble a genuine measurement (e.g., Bent et al., 2020, for wearable heart rate; Reuter et al., 2015, for MRI; Syed et al., 2020, for accelerometry). Such data bias results rather than merely adding noise (Gonsalves et al., 2026; Reuter et al., 2015). Fields that adopted sensor-based measurement well before consumer XR have not eliminated the problem, but they have established methods for managing it. Invalid data are detected with explicit rules applied to every recording (e.g., Mathôt & Vilotijević, 2023; Migueles et al., 2019), and devices’ reports on their own data have been tested against conditions in which the true state is known: when participants covered one eye on cue, two Tobii eye trackers still reported data for both eyes (Niehorster et al., 2018). Guidance in these fields recommends that such rules and their results be reported, including the quality-control checks applied, the action taken on invalid data and the amount of data excluded (Gross et al., 2013; Matthews et al., 2012; Quintana et al., 2016). Open-source tools apply such rules automatically and write a quality report for each recording or participant (Esteban et al., 2019; Migueles et al., 2019), and some store the results beside the unaltered data rather than deleting the flagged samples (Huberty et al., 2026; Lopez-Calderon & Luck, 2014).

Second, XR data-collection toolkits do not write their data in a common structure. The Brain Imaging Data Structure (BIDS; Gorgolewski et al., 2016) specifies how data are organized rather than how experiments should be designed (Jeung et al., 2024), and its EEG extension can record each channel’s status, good or bad, in a file beside the raw recording, so that such results can be exchanged between laboratories (Pernet et al., 2019). Motion-BIDS (Jeung et al., 2024) extends these conventions to motion data, a data stream recorded by XR headsets. A recent scoping review tabulates the output formats of 25 open-source VR data-collection toolkits: BIDS is not among them, although the review recommends “BIDS-like layouts where appropriate” (Thomsen & Krueger, 2026, p. 4330).

Standardizing how data are organized also presumes that the data are sound, and in XR the interface standard itself provides for data that look valid but are not current measurements. Each location that OpenXR reports carries status flags indicating whether its position and orientation contain valid data (valid) and whether they are actively tracked (tracked). When tracking is lost, the OpenXR specification states that runtimes, the software that implements OpenXR, should continue to provide inferred or last-known poses, whose position must remain marked valid and should, but need not, be marked untracked (Khronos Group, 2026; Khronos OpenXR Working Group, 2026). In preliminary testing, the Meta Quest application programming interface (API) kept Quest 2 controllers marked as tracked after they had left the view of the headset’s cameras (Costa et al., 2026). Whether a headset’s flags mark such samples therefore has to be established empirically. The XR toolkits we surveyed that record these flags do not test them under induced tracking loss, and automated checks on headset data cover either motion or gaze, not both, or produce no documented per-session quality report (see Existing tools for XR behavioral research). We found no study that reports a systematic test of whether headset status flags correctly mark induced tracking loss. Without such a test, even a correctly formatted dataset can faithfully encode unreliable measurements.

Finally, building the experiment itself remains costly. XR experiments are commonly built in game engines, software for creating interactive 3D applications, and engines such as Unity “have been created for game designers rather than behavioral scientists” (Brookes et al., 2020, p. 455). Working in them often requires specialized expertise, which places “a barrier to entry for many research groups” (Thomsen & Krueger, 2026, p. 4323). Hosseinzadeh et al. (2026), reviewing the literature on 34 VR toolboxes, identify technical overhead as the most often reported challenge: “Many toolboxes, especially those built on game engines, require a steep learning curve and proficiency in scripting languages like C#” (p. 10). Experiment builders for screen-based studies combine ease of entry with flexibility: a study can be assembled in a graphical interface with little or no programming, and code can be added where the design requires it (Henninger et al., 2022; Peirce et al., 2019). For VR toolkits, a recent review describes a recurring trade-off between ease of use and expressive power, in which researchers “often choose between fast development for standard paradigms and the engineering effort required for tailored experiments” (Thomsen & Krueger, 2026, p. 4329).

To address these three barriers, we present ResXR (Research with XR), an open-source toolkit for behavioral research with standalone Meta Quest headsets. Its Unity template records head, hand, eye and face tracking (the last two on the Meta Quest Pro) through the headset vendor’s software development kit (SDK), together with the SDK’s timestamps and status flags. Its Python pipeline, which shares no code with the template, checks each recording for hand tracking loss and eye closure, and every recorded stream for sampling-rate deviations and clock dropouts. It can mask flagged intervals in a derivative dataset, leaving the original unaltered, and exports both in Motion-BIDS format with a quality report for each session. The vendor signals that the hand-tracking-loss and eye-closure checks read were chosen by stress testing: on two Meta Quest Pro headsets, we induced tracking failures and recorded which signals marked them. The protocols are published as an open registry, so the tests can be repeated on other headsets and SDK versions. Three ready-to-run paradigms, covering discrete choice, spatial navigation and free visual exploration, let researchers collect and process data without programming. Because all experiment and data-collection code is open to modification, the paradigms also serve as starting points for tailored experiments. ResXR is hosted on GitHub, where we envision it becoming a hub for further development by the research community. An early version of ResXR was shown as a conference demonstration at IEEE VR 2026 (Bergstein, Barel, Shai Basson, et al., 2026a).

### Existing tools for XR behavioral research

Table 1 compares tools that record or process behavioral data from XR headsets. They differ in whether they keep the headset’s own report of when tracking was lost, whether they check the quality of the recorded data, and in what format they export it. This report is the set of status values the headset attaches to each sample, including the valid and tracked flags described above. Of the tools that read the headset’s interface, some store a subset of these values with every sample and others use them internally without storing them (Table 1); none tests them under induced tracking loss, and each relies on them as the runtime reports them. Our stress tests show why this matters (Meta XR Core SDK v78; Channel curation): when a hand was lost, Meta’s Body API, which reports the hands as joints of a body skeleton, moved them to a fixed arm pose and kept their valid flags set, and only their tracked flags marked the loss. LayerAwareLogging, for example, records the hands through the Body API rather than the dedicated Hand API and stores the valid flags but not the tracked flags; in our recordings, a researcher reading only the valid flags would have taken that fixed pose for the participant’s real arm position.

**Table 1.**
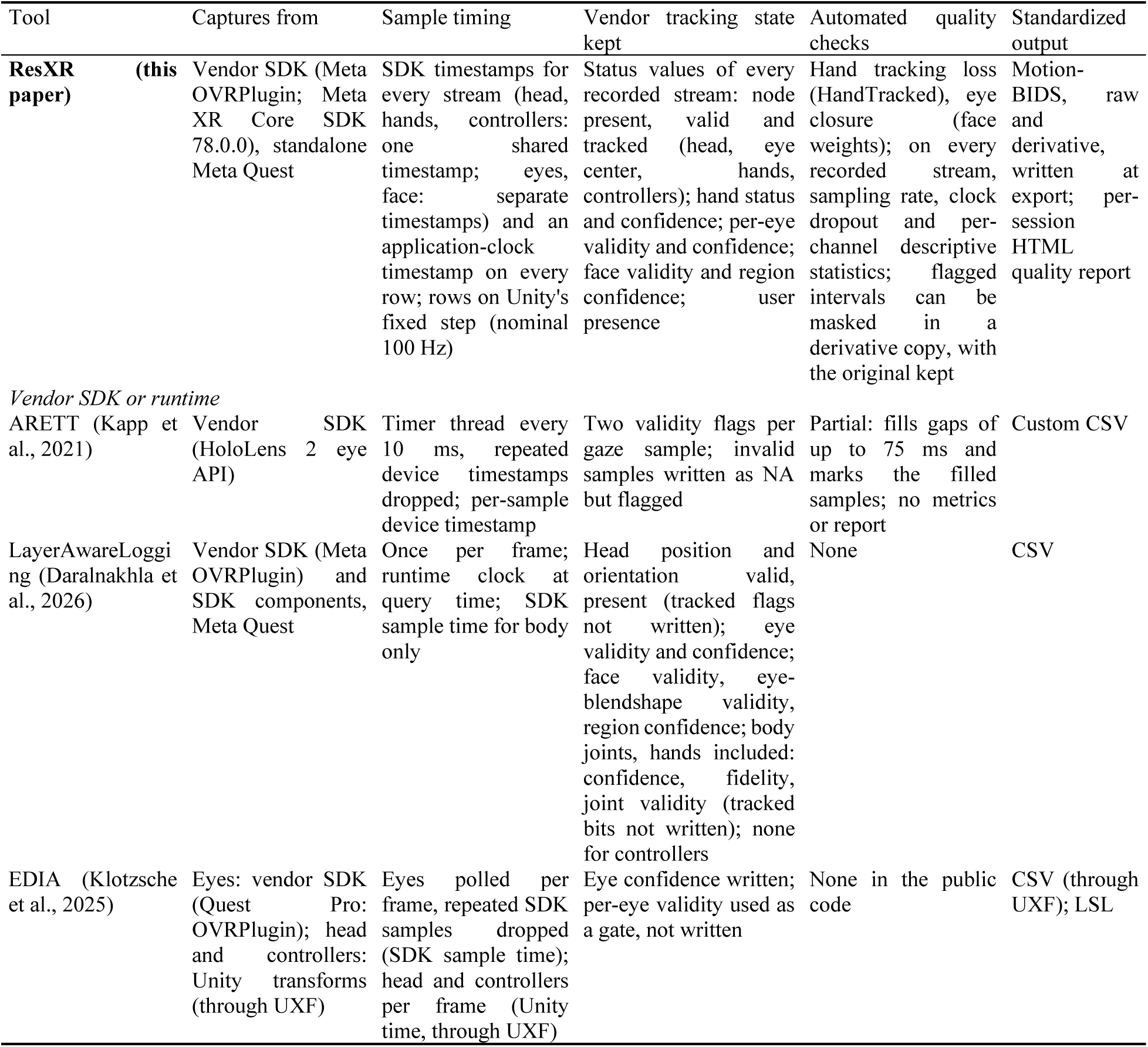
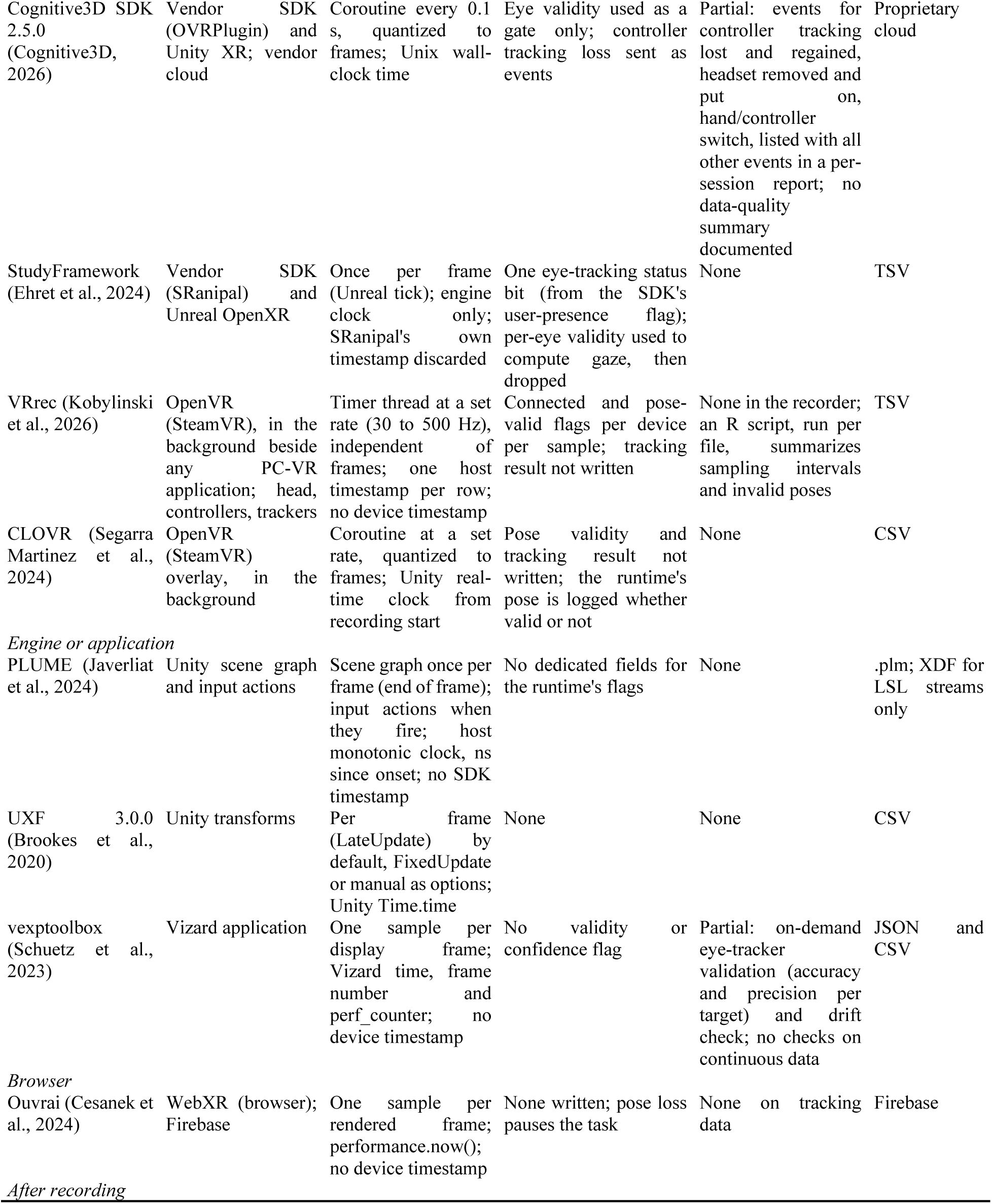
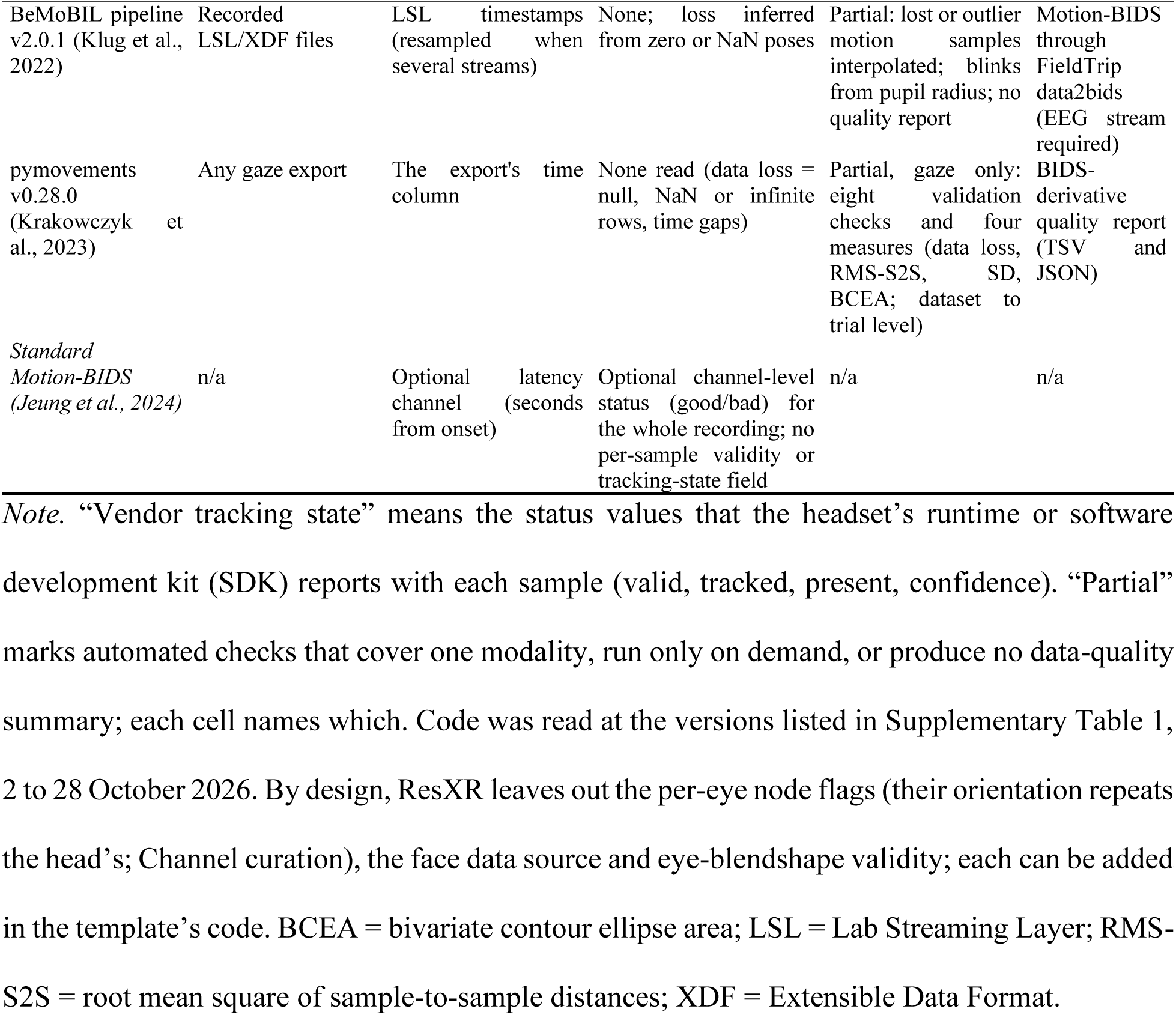
Data provenance and quality in tools that record or process behavioral data from XR headsets

Quality checks in other tools are limited to specific areas: eye-tracker accuracy and precision at validation targets, not in the continuous recording (vexptoolbox; Schuetz et al., 2023), invalid poses and sampling intervals, with a separate script run by hand (VRrec; Kobylinski et al., 2026), and lost motion samples and blinks, which are interpolated and shown in diagnostic plots without a quality report (BeMoBIL pipeline; Klug et al., 2022). None checks the full range of a headset’s tracking modalities (on the Meta Quest Pro: head, hands, controllers, eyes, face and body) in one quality report per session (Table 1). Data in Motion-BIDS can also lack the runtime’s flags: the standard defines no per-sample field for validity or tracking state (Jeung et al., 2024), and an official example dataset recorded with a headset (Gramann et al., 2024) was converted from an LSL recording and has no channels for either.

### Tool description

ResXR is an open-source toolkit that provides a complete path from immersive experiment to Motion-BIDS dataset and quality report: an extensible skeleton whose Unity template runs entirely on standalone headsets, without external hardware or a network connection, and whose Python pipeline converts those recordings into a Motion-BIDS-formatted dataset and a quality report for each session. It is organized into four functional stages (**Fig. *1***): 1) a Base Template that provides a reproducible experiment scaffold, continuously streams multimodal tracking data to storage, and records experiment-specific behavioral events in a structured, annotated format with built-in column documentation; 2) a Validation stage that evaluates data integrity through automated quality checks and flags; 3) a Preprocessing stage that can apply quality-flag-based masking to produce derivative datasets; 4) a Standardization and reporting stage that exports analysis-ready Motion-BIDS (Jeung et al., 2024) datasets, comprising sensor streams, a task-events timeline, and descriptive metadata, alongside visual quality reports. These four stages are implemented across two independent components: a Unity experiment template that covers data acquisition and experiment creation (Base Template), and a Python pipeline that covers the remaining three stages. The two components share no code; their only interface is the CSV and JSON files the template produces and the pipeline consumes. This architecture allows researchers to adopt either component independently, integrating ResXR’s data acquisition into existing analysis workflows or applying the Python pipeline to data collected outside the template once they follow the template’s file and column conventions (see Input requirements for data from other sources). The four stages address the three barriers described in the Introduction: the Base Template addresses the cost of building the experiment and records the SDK’s timestamps and status flags; the Validation and Preprocessing stages address invalid data that look valid; and the Standardization and reporting stage addresses the lack of a common structure for XR data.

**Fig. 1.**
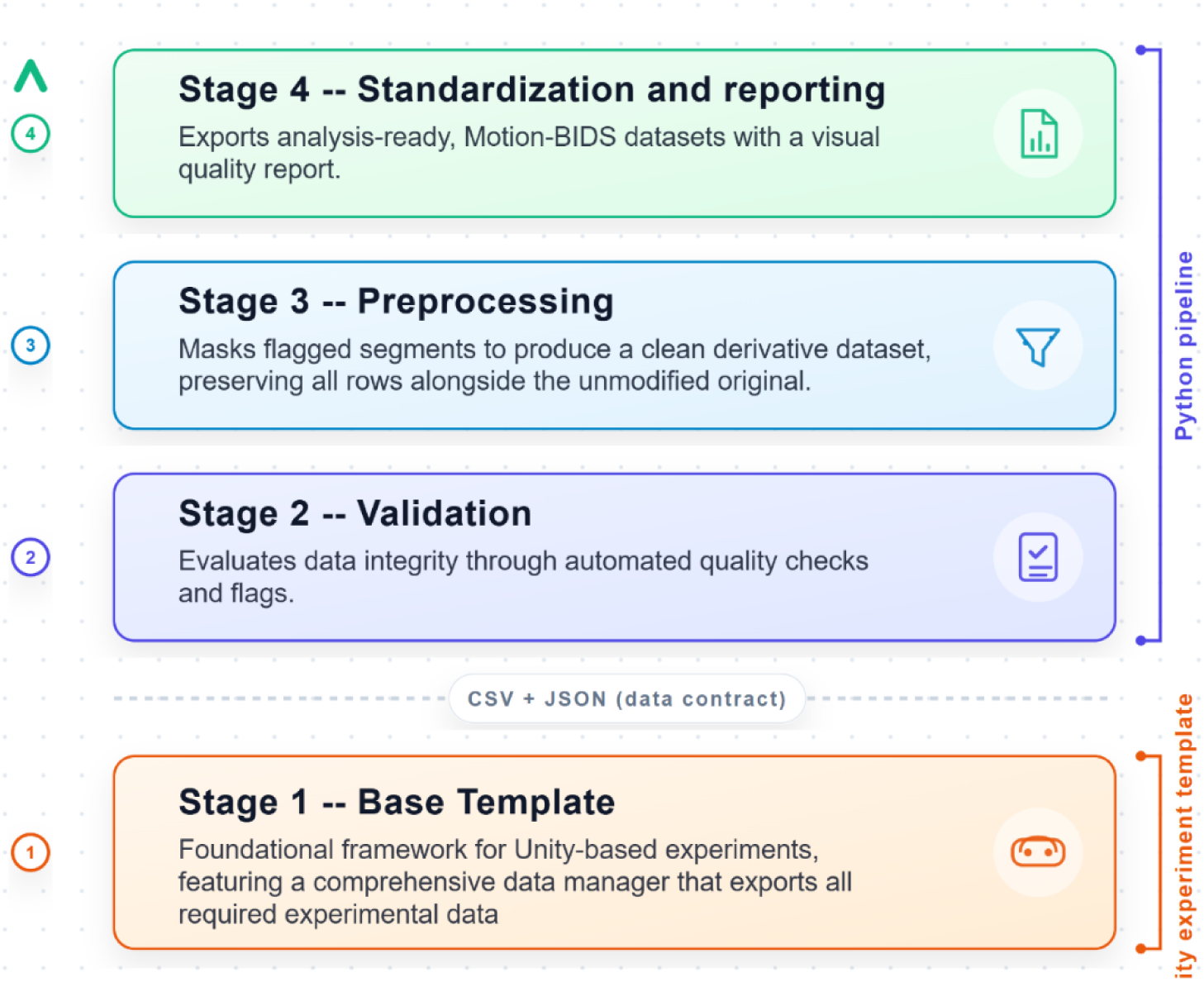
**The four-stage ResXR architecture**. Stage one (Base Template) is implemented as the Unity experiment template; stages two through four (Validation, Preprocessing, Standardization and reporting) as the Python pipeline.

### Unity experiment template

The Base Template provides a ready-to-run experiment scaffold that reduces the engineering overhead typically required to begin XR behavioral experimentation. From session onset, it continuously writes time-stamped multimodal tracking data to on-device storage. The template currently targets Meta Quest standalone headsets (Quest 2, Quest 3, Quest 3S, Quest Pro), selected on the basis of two criteria relevant to behavioral research: standalone operation, which eliminates tethering constraints that complicate deployment outside traditional laboratory settings; and native integration of multimodal sensors: the Quest Pro, the only one of these headsets with on-device eye tracking and facial expression tracking (Meta Platforms, 2026a), includes them without auxiliary hardware, reducing participant burden and hardware synchronization complexity. All recordings reported in this paper were made on Meta Quest Pro headsets.

The template employs an additive scene-loading architecture in which a persistent Base Scene, which provides the recording infrastructure, remains active throughout the experimental session (Fig. 2; Supplementary Note 5). Experiment-specific content is loaded additively on top of this persistent foundation. Because the Base Scene is never unloaded, the recorder is not destroyed or restarted during scene transitions.

**Fig. 2.**
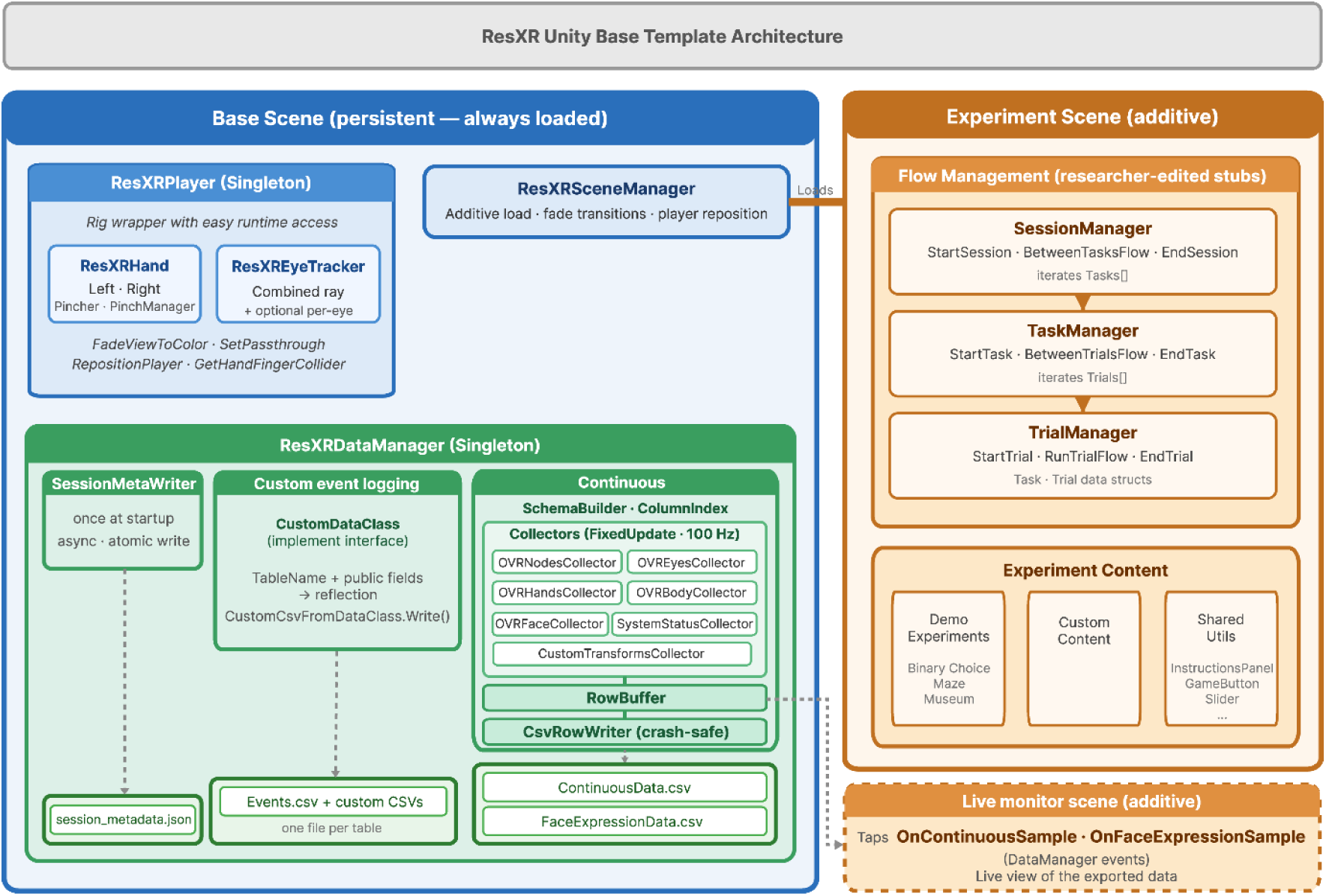
ResXR Unity Base Template architecture. The template divides into two co-loaded scenes: a persistent Base Scene that provides the recording infrastructure, and an Experiment Scene containing researcher-edited experiment flow and content. Three components occupy the Base Scene: ResXRPlayer, ResXRSceneManager and ResXRDataManager. Recorded data is written to output files including the continuous tracking CSV, face expression CSV, session metadata JSON, and researcher-defined event CSVs generated through the template’s custom data-class mechanism.

ResXRPlayer’s eye tracking sub-component, ResXREyeTracker, delivers gaze tracking, raycast-based focused object detection, and eye position data. A combined ray derived from both eyes, computed whenever both eyes’ gaze confidence is at least 0.5, records which virtual object the gaze ray hits and where (Supplementary Note 5). Complementing this scene-level gaze computation, the OVREyesCollector records per-eye gaze orientation quaternions alongside validity and confidence signals for each eye independently.

Experiments are organized through a Session → Task → Trial hierarchy implemented as three editable manager classes, following standard experimental design terminology in behavioral and cognitive science. A SessionManager iterates through tasks and handles session-level initialization; a TaskManager governs individual tasks, iterating through trials and managing transitions between trials; a TrialManager encapsulates trial-specific logic with start and end hooks. Researchers implement their experimental logic directly inside these manager classes, owning and editing them, building on the provided data logging and scene management infrastructure while maintaining full control over stimulus presentation, response collection, and trial sequencing. This three-tier structure is proposed as a default scaffold but remains subject to the researcher’s discretion: the SessionManager may be used as the sole orchestrator to accommodate any experimental flow that does not naturally decompose into tasks and trials.

Data collection follows a modular collector pattern: dedicated collector classes for each tracking modality (head, hand and controller nodes, eye gaze, hand skeleton, facial expressions, upper-body joints, system status such as user presence and recentering, and experimenter-defined transforms) are conditionally instantiated based on a per-session recording configuration. This modularity allows researchers to enable only the collectors relevant to their study, reducing file size and computational overhead through the recording configuration alone. The template focuses on the data a behavioral experiment analyzes: the participant’s tracking data as the SDK delivers it and the experiment’s events. PLUME, by contrast, was built to replay sessions with their scene context and takes eye gaze and body motion from Unity, “making it unsuitable for studies that require access to tracking data before engine-level processing” (Daralnakhla et al., 2026, p. 180). Each collector writes to a shared row buffer at every physics tick, producing a single ContinuousData.csv file in which each row represents one sampling epoch. Each row carries two classes of timestamp: a shared application-clock timestamp (Unity’s realtimeSinceStartup) that indexes the collection epoch, and the per-stream SDK timestamps described under Acquisition through the vendor SDK. Facial expression data (70 expression weights per sample) are written to a separate file to manage data volume while preserving temporal alignment through the shared application-clock timestamp. Experiment-specific behavioral events are recorded through annotated custom table classes. Each class defines onset and duration as mandatory fields, recorded in seconds on the application clock shared with the continuous data, aligning acquisition with the BIDS events model at the point of capture; in the BIDS events file, the pipeline expresses onsets relative to the start of the motion recording. Additional fields, such as reaction times, stimulus identities, and categorical responses, are annotated with a [ColumnInfo] attribute (details in Supplementary Note 1).

Several design choices address data quality at the point of collection. All enum-based sensor flags (e.g., hand tracking status, body calibration state, per-joint validity) are decomposed into individual binary columns rather than stored as string representations of the enum value, eliminating parsing ambiguity in downstream analysis and ensuring compatibility with the Python pipeline’s quality checks (see Python pipeline). The template also automatically records session metadata, including the Unity, OVRPlugin and operating-system version strings, the recording start time in UTC, the fixed timestep, enabled sensor modalities, and headset model, in a companion JSON file to support reproducibility auditing. The template also includes a Live Monitor scene that displays the recorded channels, including the validity flags, in the headset during a session.

All continuous and event data are exported as CSV files, with most continuous-data column names following a {Modality}_{Component}_{Field} convention (e.g., LeftHand_Root_px, Body_Hips_qx) to support programmatic column selection.

### Python pipeline

Following data collection with the ResXR Unity Base Template, the ResXR Python pipeline transforms raw experimental data into analysis-ready, standardized datasets. Pipeline operation requires no programming: processing settings are specified declaratively in a single configuration file, and a single command invokes the full processing chain.

The pipeline (Python ≥ 3.10) runs its stages in a fixed order coordinated by a central orchestration module (Fig. 3). The configuration file, in YAML, specifies input paths, device metadata, tracking system toggles (head, hands, eyes, face, body and controllers; the last two are off in the shipped configuration), expected sampling frequencies, quality-check parameters, and BIDS output conventions. Researchers interact with the pipeline through either a command-line interface or a programmatic Python API, both of which invoke the same orchestration logic.

**Fig. 3.**
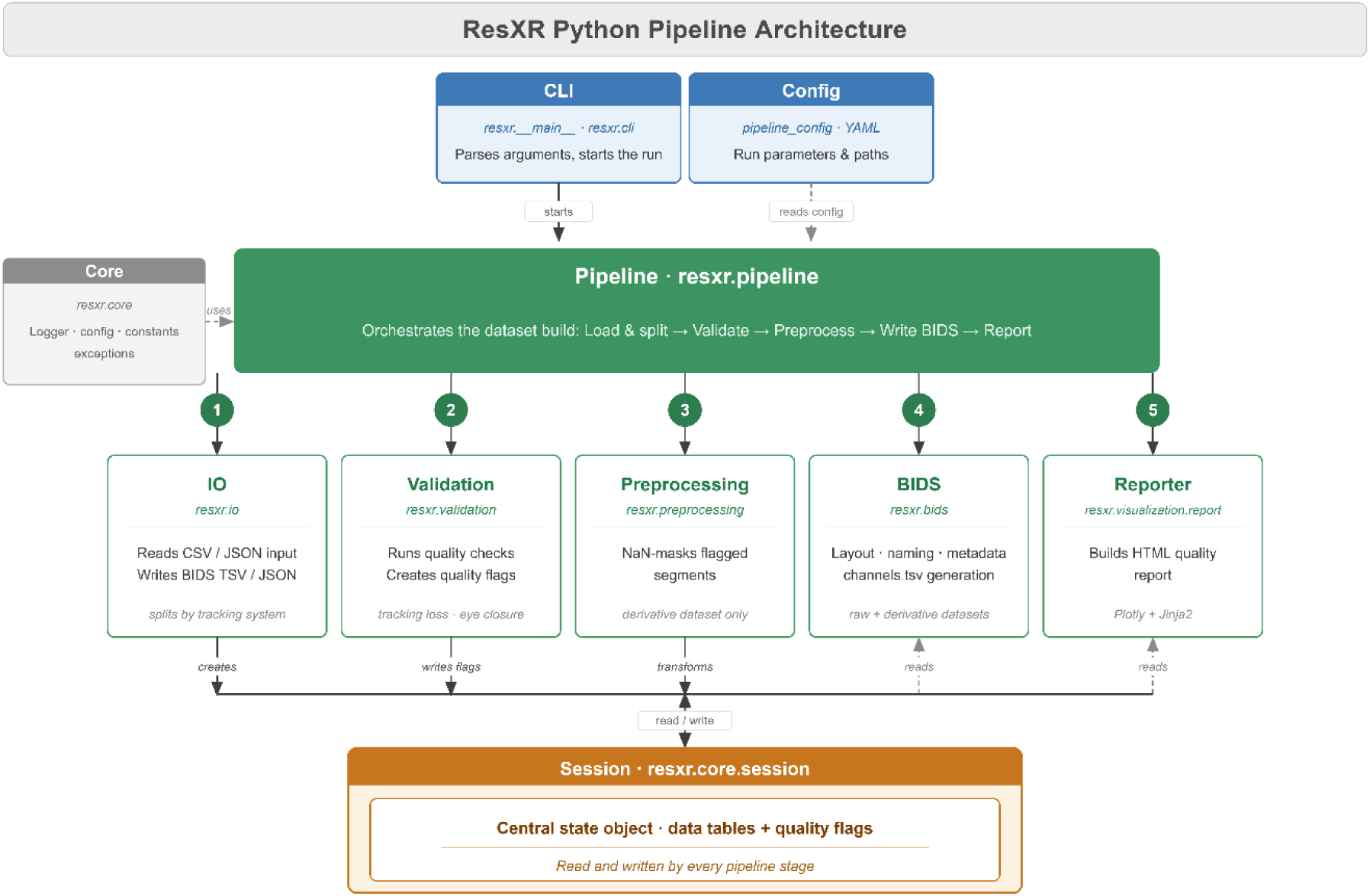
ResXR Python pipeline architecture. A single orchestrator (resxr.pipeline) runs its stages in fixed order on input exported by the Unity template: IO reads the raw CSV/JSON and custom table files and splits the continuous file into per-system streams; Validation flags quality issues (e.g., hand tracking loss, eye closure); the pipeline constructs the unified task-events timeline; BIDS writes the raw Motion-BIDS dataset; Preprocessing can NaN-mask flagged segments to produce the derivative data, leaving the raw dataset intact; BIDS writes the derivative dataset; and Reporter renders an HTML quality report. Solid arrows denote writes; dashed arrows denote reads and dependencies.

#### Validation

Five quality checks are provided in pipeline v0.2.2, and researchers can add their own and choose in the configuration file which checks run (Supplementary Note 4): hand tracking loss detection, which reads the HandTracked status fields selected by the stress tests (Evaluation); sampling rate consistency, which raises a flag when a stream’s effective sampling frequency deviates from the expected rate by more than a configurable tolerance, and a second flag when the intervals between the stream’s own samples, read from its SDK timestamps, are irregular; clock dropout detection, which flags per-system timestamps that drop to zero or are missing during the recording; eye closure detection, a cross-stream check that flags intervals during which both face-expression blend-shape weights (Eyes_Closed_L and Eyes_Closed_R) reach or exceed a threshold for at least a minimum duration, then propagates the resulting flags from the face stream to the eye-tracking stream for optional downstream filtering; and per-column descriptive statistics (count, mean, median, standard deviation, minimum, maximum, 5th/25th/75th/95th percentiles, and NaN proportion) for all channels in each stream. Default thresholds, the check interface and the flag record are described in Supplementary Note 4.

#### Preprocessing

The pipeline can apply quality-flag masking to produce a derivative dataset; masking is off by default. Flagged time segments are replaced with NaN values rather than deleted, preserving row count and temporal continuity consistent with BIDS conventions. Masking operates at column-level granularity: a hand-tracking-loss flag masks that hand’s position, orientation, scale and confidence columns and keeps its validity, status and timing columns, and an eye-closure flag masks the gaze columns of the eye-tracking stream and leaves the face stream unchanged. When masking is enabled, flags from hand tracking loss, clock dropout and eye closure detection contribute by default; sampling rate flags are advisory only and never contribute to masking. The researcher controls which checks contribute to masking through the configuration file; with masking off, the derivative dataset’s motion and channel files are identical to those of the raw dataset, and the flags appear in the quality report. The raw dataset is never modified.

### Standardization and reporting

The pipeline writes each session into two BIDS dataset tiers: a raw dataset containing the original data and a derivative dataset (masked only when masking is enabled; see above). For each tracking system, the exporter generates four files: a headerless motion.tsv file whose column semantics are defined in the accompanying channels.tsv, and channels.json and motion.json sidecars (Supplementary Note 4).

By default, the pipeline generates an HTML report for each session synthesizing quality-check results into a single document. Reports include a session header with device and acquisition metadata, a summary (duration and the number of warning-level flags), an interactive timeline visualization (built with Plotly (Plotly Technologies Inc., 2015)) displaying flagged segments and optional event markers on a shared time axis, per-stream statistics tables (row counts, channel counts, NaN percentages, expected versus effective sampling rates), and a quality flags table with all flag times expressed in global time relative to recording onset. Each report also records when it was generated and the pipeline version that produced it. Each report is a single self-contained HTML file that opens in any web browser without a network connection and can be shared alongside the BIDS dataset for peer review of data quality decisions (Fig. 4).

**Fig. 4.**
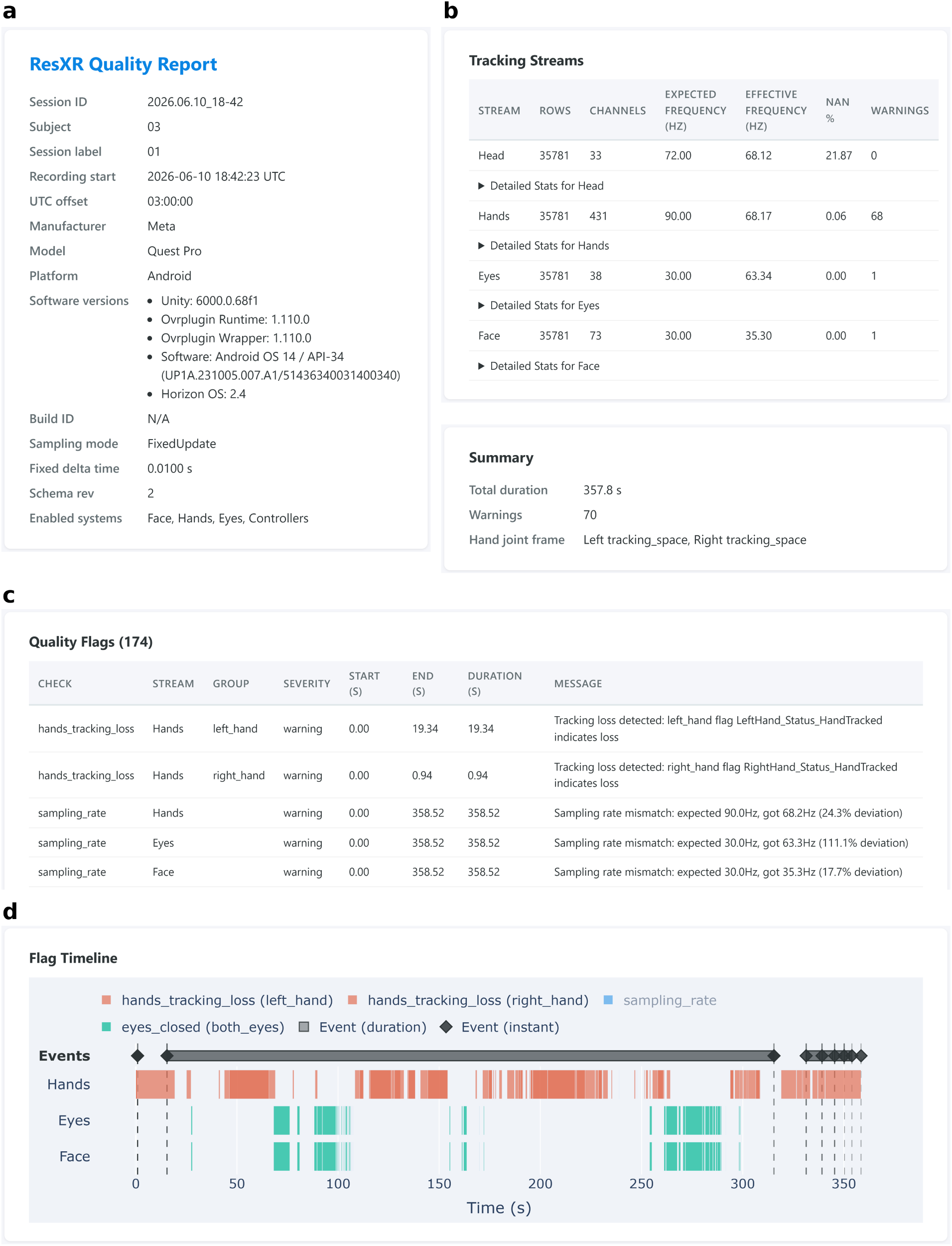
Self-contained HTML quality report for a representative ResXR session. A single self-contained HTML file summarizing data quality for one 357.8 s Museum Viewing recording from a Meta Quest Pro (subject sub-03 of the demonstration dataset), shown here as its four constituent panels: (a) session header and summary, (b) per-stream statistics, (c) quality-flag table (n = 174 for this session) and (d) flag timeline, with all flags on one shared time axis (seconds from recording onset), color-coded by check and stream. Panel contents are listed in Supplementary Note 4.

### Input requirements for data from other sources

The pipeline also accepts data recorded without the template, provided that the files follow the template’s conventions. Each session is a folder with a continuous-data CSV file and a JSON metadata file (the file-name patterns are configurable); face, event and custom-table files are optional. The continuous file must contain a timeSinceStartup column in seconds, which defines the session clock, and event onsets must use the same clock. Columns are assigned to tracking systems by their name prefix (e.g., LeftHand_; Supplementary Table 2), and channel types and units are inferred from the name suffix (e.g., _px, a position in meters). The metadata file states which systems were recorded. A session that lacks a required file or any valid timeSinceStartup value is skipped with an error. Supplementary Note 4 gives the full contract and the behavior for departures from it; data that meet it pass through the same quality checks and are exported to the same Motion-BIDS dataset and report as data recorded with the template.

### Adding a sensor modality

New sensor streams are added by editing each component’s code. In the template, a new stream is a collector class that is registered with the data manager and whose columns are declared in the schema (ResXR Core Team, n.d.); in the pipeline, it is a new tracking system defined by its column prefixes, after which it is split, checked and exported like the existing systems (Supplementary Note 4). Signals from external devices can be stored in the same BIDS dataset in the format the specification defines for their modality, for example cardiac recordings, EEG or electromyography (BIDS Contributors, 2026); ResXR neither writes nor checks them, and the researcher must align their clock with the headset’s, for example from an event recorded by both devices.

## Design rationale

### Acquisition through the vendor SDK

At the software level, the Unity template is built on Unity 6 and the Meta XR Core SDK v78 (OVRPlugin 1.110.0). It reads sensor data through OVRPlugin, the native plug-in through which Meta’s Unity SDK queries the headset runtime, rather than through Meta’s higher-level C# components (such as OVRHand) or Unity’s cross-vendor OpenXR input features. The exception is the gaze-hit record: ResXREyeTracker casts its rays from the eye transforms that Meta’s OVREyeGaze component updates (see Unity experiment template). While OpenXR provides multi-platform compatibility, neither its core specification nor any multi-vendor extension includes per-eye gaze orientation or facial expression weights; on Meta headsets, OpenXR offers them only through Meta’s own extensions (Khronos OpenXR Working Group, 2026). OVRPlugin exposes every tracked modality, together with its status fields, through one interface. Direct OVRPlugin access is further motivated by a critical requirement for temporal precision: it exposes per-sample SDK timestamps (head and hands: one shared timestamp; eyes and face: separate timestamps) in addition to the application-loop timestamp. Application-loop timestamps record when the application queried the SDK. Because the logger runs faster than the SDK delivers new values, successive queries return the same value, and application-loop timestamps give each repeat a distinct time. SDK timestamps make these repeats explicit: rows that repeat a value share its timestamp (see Fixed-rate logging), so analysts can identify them. This distinction matters both for relating each recorded row to the SDK value it holds and for multimodal temporal alignment across streams.

### Inspectable and modifiable code

The Unity template is designed as a glass-box framework, a term we borrow from fMRIPrep (Esteban et al., 2019): rather than providing an opaque library consumed through a fixed API, it exposes all experiment and data-collection code for direct inspection and modification by the researcher. Base classes, flow-management logic, and sensor-access routines ship as source files, to be inspected and modified directly, so that the final experiment code is fully owned and understood by the research team.

### Fixed-rate logging

Unity runs scripts in two loops: Update, the main frame update, which runs once for every displayed frame, and FixedUpdate, which runs at a fixed rate set in the project (Unity Technologies, n.d.). Unity’s default execution logic, built on Update, is “intrinsically coupled to the rendering loop,” leaving “data logging susceptible to framerate-induced variability” (Hosseinzadeh et al., 2026, p. 12). The template therefore logs every stream in FixedUpdate rather than Update, at 100 times per second, so the number of rows depends only on the duration of a recording, not on the rendering load. This rate is higher than the rate at which the runtime delivers new head and hand poses, so some rows repeat the previous one, and the SDK timestamp in each row marks which rows hold a new sample (see Processing the example sessions). We chose this rate to exceed the headset’s frame rate (about 72 frames per second; see Performance cost of continuous logging). At a lower fixed rate, Unity would skip FixedUpdate in some frames, and the tracking data of those frames would be lost, indistinguishable in the file from data the headset never sent. Above the frame rate, Unity runs FixedUpdate at least once in every frame (Unity Technologies, n.d.), so the tracking data of every frame reach the file.

### Plain-text output

The Unity template writes its continuous streams as plain-text CSV. This is a deliberate trade against binary serialization, and the main reason is that a plain-text record can be opened in a text editor and read directly into standard analysis workflows in Python, R and MATLAB, without the software that wrote it or a separate reader library; CSV, valued for this integration with analysis software, is the most common export format among VR research toolboxes (Hosseinzadeh et al., 2026, p. 8). For data intended to be shared, reanalyzed and read years after collection, that independence is itself a reproducibility property; BIDS likewise stores metadata and tabular data as text (Gorgolewski et al., 2016). A second reason is the archival format: Motion-BIDS, the standard the pipeline targets, specifies tab-separated text for motion data, channel descriptors and event timelines (Jeung et al., 2024; Supplementary Note 5).

After every row, the data are passed to the headset’s operating system, which writes them to storage. A session interrupted by an application crash, headset removal, or experimenter termination therefore keeps every complete row recorded up to that point, losing at most the row being written. The cost of these choices was measured (see Performance cost of continuous logging). In practice the cost was small: with logging on, the frame rate was unchanged, and the main effect was about one additional repeated frame per second. Storage is not a constraint either: the default configuration writes about 1.46 GB per hour of recording, so a hundred one-hour sessions occupy about 146 GB, 57% of the 256 GB of a Meta Quest Pro (Meta Platforms, 2022).

### Example use

To reduce the time from installation to initial data collection, the template includes three ready-to-use experiment implementations spanning common behavioral research designs (Fig. 5). Each follows the Session → Task → Trial structure described above. The same Base Template supports direct use, adaptation, and entirely new paradigms. These three paradigms were selected to cover a representative range of experimental constraints and tracking modalities: structured discrete responses with hand-tracked input (*Binary Choice*), goal-directed continuous spatial navigation (*Maze Navigation*), and free visual exploration with eye-tracked gaze (Museum Viewing). Together they exercise the template’s core capabilities, including trial-based event logging, continuous spatial tracking, and gaze-ray hits on virtual objects, while demonstrating that the Session → Task → Trial hierarchy accommodates designs of varying complexity without modification to the underlying data collection infrastructure. The paradigms share reusable components (instruction panels, buttons, a rating slider and start-position markers) that new experiments can use.

**Fig. 5.**
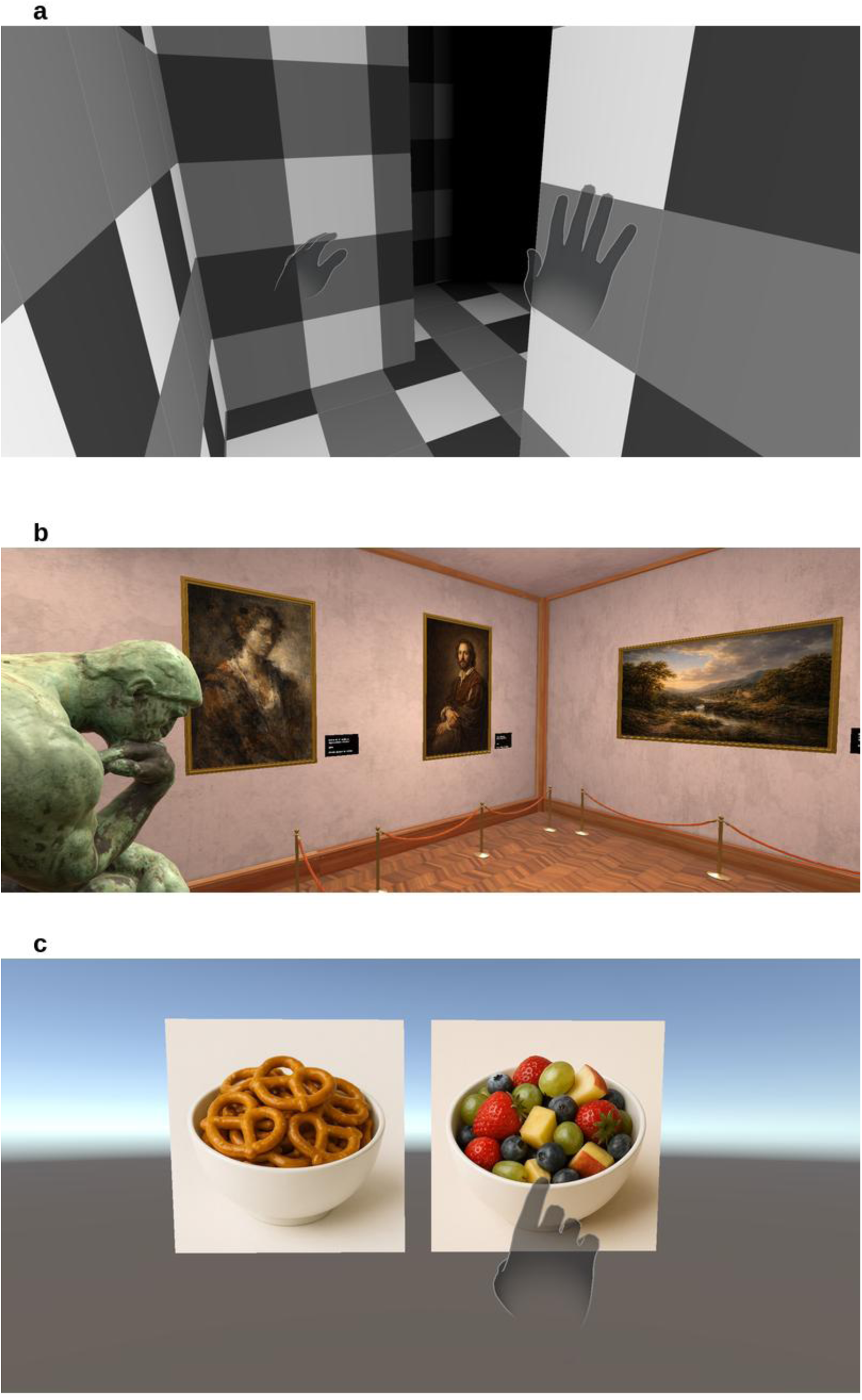
**ResXR demonstration paradigms**. Three ready-to-run experiments are included with the ResXR Unity template, each built entirely on the template’s standard components, exactly as a researcher would build their own experiment. (a) Maze Navigation: participants navigate a maze to collect a golden coin. (b) Museum Viewing: participants freely explore a virtual gallery and view artworks. (c) Binary Choice: participants choose between the two stimuli of a simultaneously presented pair.

### Discrete choice

A *Binary Choice* paradigm implements a two-alternative forced-choice (2AFC) design: stimulus pairs are loaded from a configurable directory in fixed or random order, presented simultaneously, and selected via hand-tracked touch input. The system records choice identity, the hand used and reaction time as trial-level events alongside the continuous multimodal data stream.

### Spatial navigation

A *Maze Navigation* paradigm presents experimenter-defined spatial layouts in which participants navigate to collect target objects, capturing continuous movement trajectories, coin-collection timestamps, start-zone entries, and a mid-task maze-rotation manipulation for analysis of spatial navigation strategies.

### Free visual exploration

A *Museum Viewing* paradigm places participants in a virtual gallery for free exploration of artwork displays, recording continuous gaze and the artwork hit by the template’s combined gaze ray, together with each artwork’s spatial bounds for offline gaze analysis and movement patterns throughout the environment, followed by a structured task in which participants rate each image on a virtual slider.

### Processing the example sessions

All three paradigms have been implemented using the annotated custom table format described in Tool description. Behavioral outcomes, trial-level choices, reaction times, and stimulus geometry in Binary Choice; per-trial maze layouts and outcomes in Maze Navigation; and artwork geometry, image ratings, and rating-scale configuration in Museum Viewing appear in the BIDS events timeline as onset-ordered rows, merged with natively logged task events such as trial onsets, coin pickups, and task transitions, with each behavioral column’s description, format, and units derived from the field annotations in the experiment code and carried through to the events sidecar unaltered (Supplementary Note 1). Extending any paradigm with a new behavioral variable requires only annotating the corresponding field with a description and a format; both the data and its documentation then propagate automatically into the standardized output without pipeline changes.

To demonstrate pipeline operation, we processed one recorded session from each of the three demonstration paradigms, all acquired on a Meta Quest Pro in standalone mode (OVRPlugin 1.110.0, Unity 6000.0.68f1) with four active tracking systems (Head, Hands, Eyes, and Face): a 35.6 s Binary Choice session, a 104.7 s Maze Navigation session, and a 357.8 s Museum Viewing session. A single configuration file mapped the three source directories to subjects sub-01 through sub-03, and a single pipeline invocation produced one BIDS dataset containing raw and derivative tiers for all three sessions, with 583 channels per session (Head 35, Hands 433, Eyes 40, Face 75, including per-stream latency channels) and a self-contained HTML quality report for each session (Fig. 4). Each session’s event timeline was written as events.tsv at the session root, with an events.json sidecar generated from the acquisition schema documenting onset, duration, and all behavioral columns with their description, units, and categorical levels. The quality checks raised 228 flags across the three sessions: 104 for hand tracking loss, 12 for sampling rate and 112 for eye closure (56 flag episodes, each flagged on the Face and the Eyes stream); the clock-dropout check raised none. The sampling-rate flags are advisory: eight come from comparing the measured rates given below with the expected rates of the shipped configuration (Hands 90 Hz, Eyes 30 Hz, Face 30 Hz), and four mark irregular sampling in the Maze Navigation session, whose recording includes pauses of up to 2 s; researchers can set the expected rates and tolerances in the configuration file. The three demonstration sessions, the configuration file, and the resulting BIDS output are distributed with release v0.2.2 of the project repository (https://github.com/ResXR/resxr-python-pipeline).

Effective rates differed by modality (head 67–71 Hz, hands 67–72 Hz, eyes 57–66 Hz, face 31–36 Hz across the three sessions; Supplementary Table 3), measured on the demonstration stack (Horizon OS 2.4, OVRPlugin 1.110.0). For head and hands, these are the rates at which their shared SDK timestamp advanced, not necessarily sensor sampling rates. Because these rates were measured on one software stack, they should be re-measured after updates, and because the streams are asynchronous, cross-modal analyses require interpolation or alignment.

## Evaluation

### Validity-flag audit under induced tracking failure

#### Procedure

The Meta XR Core SDK provides access to six sensor modality groups (head, eyes, hands, face, body, and controllers) through multiple overlapping APIs. Depending on the modality, these APIs expose kinematic data (position and orientation), gaze orientation quaternions, or facial expression weights, each accompanied by internal state variables such as per-sample validity flags, tracking status indicators, and confidence scores. Because the pipeline’s hand-tracking-loss and eye-closure checks read signals selected from these channels, we conducted a systematic empirical evaluation to determine which channels produce reliable, non-redundant data and which internal state variables accurately distinguish measured from inferred tracking states.

To evaluate whether the extracted validity flags and confidence scores accurately reflect sensor state, we designed 17 controlled stress-test protocols (Supplementary Table 4); ten annotated sessions (eleven recordings) were recorded in a dedicated XR testing arena using Meta Quest Pro headsets across several template builds. Each test session followed a choreographed protocol designed to trigger specific tracking failure modes, generating datasets containing both intervals of maintained tracking and intentional signal interruptions at annotated timepoints.

For hand tracking, protocols tested the boundaries of the headset’s inside-out camera field of view, included bilateral arm swings and introduced graded occlusion; for eye tracking, they included gaze shifts to cardinal positions, gaze fixation during head rotation, monocular occlusion, single-eye closure and sustained eye closure; for face tracking, sustained expressions, partial occlusion of the lower face and headset removal (Supplementary Table 4; Supplementary Note 2).

We synchronized each test session’s data with a ground-truth event timeline in which a researcher performing each choreographed protocol noted the timestamps of intentional tracking manipulations (Supplementary Note 2). Because event boundaries were annotated via visual inspection rather than hardware-synchronized triggers, temporal precision of the ground-truth timeline is on the order of seconds rather than milliseconds; analysis therefore focused on whether flags exhibited correct state transitions within the annotated event windows rather than on sub-second transition latencies. Evaluation criteria were temporal concordance (whether each flag transitioned during the corresponding annotated event window) and cross-API consistency (whether redundant signals across the Node, dedicated Hand and Eye, and Body APIs agreed on tracking state and pose).

The full set of stress-test protocols is provided in Supplementary Table 4 and is maintained as a community registry (Bergstein, Barel, Shai Basson, et al., 2026c) at https://github.com/ResXR/xr-validation-protocols (release v1.1.0, https://doi.org/10.5281/zenodo.23679224). The validity-flag audit was conducted on two Meta Quest Pro headsets (firmware v79 and v83 where the laboratory’s test notes record it), with Meta XR Core SDK v78 (OVRPlugin 1.110.0) and Unity 6000.0.40f1 or 6000.0.60f1; the demonstration sessions (Example use) were recorded later with the same OVRPlugin version, Unity 6000.0.68f1 and Horizon OS 2.4, and include no induced tracking failures.

#### Channel curation

The column registry (Supplementary Data 1; procedure in Supplementary Note 2) lists 1,460 candidate columns across six modality groups; 1,436 of them from OVRPlugin. OVRPlugin’s overlapping APIs proved substantially redundant: independent interfaces reported the same hand positions in every hand-tracked row, identical in all seven significant digits the template writes; several fields, including the hand nodes’ velocity, were not populated by Meta XR Core SDK v78 (OVRPlugin 1.110.0).

The Body API was excluded from the default schema. On Meta Quest Pro, body joints are not directly measured but inferred from headset, hand and controller movements (Meta Platforms, 2026a). During induced hand-tracking loss, the Body API’s hand and arm joints did not drop to zero, as the Hand API’s positions did, but moved to a fixed arm pose while remaining flagged as valid; the Body API’s confidence and calibration status did not change, and only the tracked bits of the lost side’s hand and arm joints marked the loss. While the hands were tracked, the Body API’s wrist agreed closely with the Hand API’s. Node-level eye quaternions reflected head orientation, not gaze, so the dedicated Eye API was retained. Curation reduced the 1,460 columns to a 607-column default schema, with face-expression weights stored separately.

#### Flag behavior by modality

Empirical stress testing showed that the SDK’s Valid and Tracked flags frequently diverge from what their names imply, with flags within one API behaving asymmetrically, so names alone risk treating inferred or stale data as genuine measurements.

### Hand tracking

Within a single API, Valid_Orientation stayed set during tracking loss while Valid_Position dropped to zero; position is therefore the reliable indicator, and HandTracked (dedicated Hand API) dropped at every induced tracking loss. Per-finger confidence did not mark finger occlusion and should not be read as a finger-occlusion signal.

### Eye tracking

Eye-tracking validity flags stayed at 1 throughout the induced failures, under monocular occlusion, eye closure and gaze-limit conditions alike, providing no usable signal; eye closure was instead marked by the face-expression weights (Eyes_Closed_L, Eyes_Closed_R), which should be used for eye-closure quality control, although how much of each closure they mark depends on the detection threshold.

### Face tracking

The Face_Status flag, the eye validity flags and the head node’s valid and tracked flags all stayed set while the headset was off. None of the signals recorded in the stress tests unambiguously distinguishes headset removal from other interruptions. Studies requiring continuous presence verification must detect headset-off at the protocol level.

### Trusted flag selection

Across the four modalities, hand tracking produced vendor signals that responded reliably to loss. Eye and face tracking required secondary signals, and no signal recorded in the stress tests distinguished headset removal from other interruptions. HandTracked serves as the primary hand tracking indicator, eye-closure detection uses face-expression eye-close weights, and headset-off detection is handled at the protocol level. Each trusted flag maps to the columns it governs for column-level masking (Supplementary Table 5).

### Performance cost of continuous logging

Recording every sensor stream 100 times per second could slow the headset and disturb the experiment. We tested this on one Meta Quest Pro running the Maze Navigation demonstration, with two runs that recorded the default schema and two baseline runs that switched the continuous collectors off and recorded only the face data and a timestamp (Supplementary Note 6). Two tools measured the cost. A log written by the application once per second recorded how much of the application’s main-thread time the logger used, and OVR Metrics Tool (Meta Platforms, 2026b), Meta’s on-device performance monitor, recorded frame timing, repeated frames, tracking latency and power once per second in one run of each condition. Its tracking measures (among them frame-to-frame, processing and overall latency) are named in its export but not defined in Meta’s documentation.

Continuous logging did not change the frame rate (72.1 frames per second with logging, 71.9 without), and no rows were lost during the task. Its cost appeared in the slowest frames. The 95th-percentile frame time, which the slowest 5% of frames in each second exceed, was 17.4 ms with logging and 15.7 to 16.2 ms without, against the 13.9 ms available per frame, and repeated (stale) frames rose from 0.65 to 1.68 per second. The logger used about 1.6 ms of the application’s main-thread time per 100 Hz tick: two thirds of it to write the rows as text, and about 5% to hand the rows to the operating system for writing to storage. The hand, face and eye frame-to-frame latencies that the tool reports differed between conditions by less than 4%. Face processing and overall latency, eye overall latency and hand processing latency differed between conditions by less than 1 ms in either direction, but hand overall latency was 12 ms higher in the run with logging. Device power was 1.1% higher with logging, and the headset’s power level, which Meta’s documentation uses to indicate when heat affects performance, stayed at normal in both runs (Meta Platforms, 2026d).

Overall, we found that continuous logging did not disrupt the experiment. With every default stream recorded at 100 Hz, the headset kept its frame rate of about 72 frames per second, every row of the task was written, and its power level stayed normal; the cost was confined to the slowest frames, about one more repeated frame per second. The 100 Hz rate is a project setting (Unity’s fixed timestep, which also sets the physics update rate). Because head and hand poses were renewed 67 to 72 times per second in the demonstration sessions, a researcher can lower the rate toward that value, so that fewer rows repeat the previous one and the logger has less to write.

### Conformance to Motion-BIDS

Both dataset tiers (raw and derivative), each containing all three demonstration sessions, were checked with the BIDS Validator (version 3.0.2) (Appelhoff et al., 2026), which reported no errors and only warnings, about recommended metadata fields, the number of listed authors and the length of the README.

## Discussion

Behavioral XR research has lacked an accessible experiment template and shared data conventions. Even studies built on identical hardware produce incompatible outputs, because no shared convention governs what to record or how to check it. ResXR addresses this gap with an experiment template, automated quality checks on the recorded data, and Motion-BIDS output that passed the official BIDS Validator without errors.

ResXR brings together, in one open toolkit, what behavioral XR studies have so far had to build separately. Three paradigms, for discrete choice, spatial navigation and free visual exploration, run out of the box, so researchers can collect data without programming. A single configuration file and a single command turn each recording into a Motion-BIDS dataset. Its events file carries the column descriptions written in the experiment code, and its quality report can be shared with the data. The stress tests supply the systematic test of headset status flags under induced tracking loss that we found missing in earlier work, and they determine which signals ResXR’s quality checks read. Recording every default stream at 100 Hz did not disrupt the experiment: the headset’s frame rate was unchanged.

Under induced tracking failure, several of the status flags described in the Introduction did not mark the failure as their names suggest (Meta XR Core SDK v78, OVRPlugin 1.110.0, on Meta Quest Pro headsets with firmware v79 and v83), making some of them an unreliable basis for quality control. In the stress tests, we identified five classes of this misbehavior: the hand Valid_Orientation bit stayed set while hand position was lost, the Body API’s arm joints moved to a fixed pose while remaining flagged as valid, per-finger confidence did not mark finger occlusion, the eye-tracking validity flags stayed set during occlusion and eye closure, and none of the signals recorded unambiguously distinguished headset removal (Evaluation). To our knowledge, these flag behaviors have not been systematically documented in the XR research literature. A researcher who encounters flags named “Valid” and “Tracked” will reasonably assume they indicate data quality, and filter accordingly. Our findings show that assumption is not entirely reliable. Empirical stress testing identified a trusted subset (HandTracked for hand tracking loss; the face-expression eye-closure weights for eye closures, whose coverage of each closure depends on the detection threshold), but which flags to trust cannot be determined from the SDK’s documentation alone.

ResXR turns these findings into defaults. The template records every stream through the vendor SDK and keeps each sample’s SDK timestamp and status flags. The pipeline flags hand tracking loss from HandTracked, which dropped at every induced loss. It flags eye closure from the face-stream Eyes_Closed weights, because the eye validity flags did not mark closure. The same scaffold ran three different designs, discrete choice, spatial navigation and free visual exploration, without changes to the recording code, which suggests that it accommodates major families of behavioral designs.

ResXR complements existing XR research tools rather than replacing them: ResXR’s data layer could be added to a UXF experiment (Brookes et al., 2020), although this combination has not been tested. The Unity template is a starting project for a new experiment, not a package that can be installed into an existing Unity project.

ResXR is designed to grow with its users. Each paradigm is a working example that researchers can copy and adapt, and new paradigms built on the template can be contributed back to the repository. The stress-test protocols form an open registry, so laboratories can audit their own headsets and SDK versions and add the results, extending the set of trusted flags beyond the Quest Pro. New quality checks can be added to the pipeline through its check interface (Supplementary Note 4). Every study built on ResXR can thus add to a shared, tested foundation for behavioral XR research.

## Scope and limitations

ResXR’s current scope is primarily Meta standalone hardware accessed through OVRPlugin, covering head, hand, eye, and face tracking. It is the configuration in which the validity-flag audit was carried out and a trusted subset empirically established. Among Quest headsets, eye tracking and camera-based face tracking are available only on the Quest Pro (Meta Platforms, 2026a), whose face weights the eye-closure check reads. The Quest Pro is no longer sold (Hayden, 2025). The template’s build settings also target the Quest 2, Quest 3 and Quest 3S, whose hand-tracking flags have not yet been audited. Five methodological constraints follow. Validity-flag behavior was characterized on one SDK version and one headset model and may differ in other releases or Quest models, so it should not be assumed to transfer without re-testing with the published protocols; the stress tests themselves involved one wearer per session, two headsets and few performances of each action per session, with ground truth annotated in whole seconds, so they support counts of detected and missed events but no false-alarm rate per unit time. The audit characterizes validity flags, not pose-level accuracy against ground truth; nor does ResXR measure the spatial accuracy or precision of gaze, which GazeMetrics measures for head-mounted eye trackers (Adhanom et al., 2020). Motion-to-photon latency was not measured, because Meta’s Unity SDK does not report it. All characterizations used standalone operation; in three brief tethered PCVR sessions (Supplementary Note 3) the Unity template recorded head, hand, eye and face tracking, but systematic PCVR testing was not conducted. The pipeline also does not test for repeated or held samples: for head and hands the SDK timestamp is one shared value that keeps advancing with the head, so a timestamp check would not reveal a hand pose that the runtime holds unchanged. The position embedded in ResXR is that empirical validation, not hardware reach, defines what the data are good for.

Tracking data raise privacy questions that the toolkit leaves to the researcher. Eye-tracking and facial-expression data can reveal sensitive information about the user, including identity (Daralnakhla et al., 2026; Jakobi et al., 2024), and head and hand tracking data can identify individual users (Bailenson et al., 2025). ResXR records to the headset’s storage without a network connection, but its Motion-BIDS output contains the raw eye and face streams; before sharing, researchers should obtain consent that covers these data and decide whether to remove or aggregate them (Thomsen & Krueger, 2026), as with other identifiable data in BIDS datasets. The pipeline copies every raw session file unchanged into the dataset’s sourcedata folder, including streams switched off in the pipeline’s configuration.

## Future directions

The architecture is designed to extend to future XR devices, including full-face headsets and smart glasses (Kim & Choi, 2021; Kugler, 2025; Mansour et al., 2025): the Python pipeline accepts recordings from any logger that writes the template’s file and column conventions (see Input requirements for data from other sources). Two extensions follow: a hardware-abstraction layer for additional XR headsets, and physiological co-registration with EEG, ECG, and fNIRS streams under BIDS conventions. Meta VR Glasses, announced in September 2026, run the same operating system and software development kits as Quest headsets (Meta Platforms, 2026c); because the template reads tracking data through Meta’s SDK, they are the closest candidate for extending it, once the SDK is updated and the status flags are re-audited on that device.

ResXR is an attempt to provide scaffolding for reproducible XR experimentation with current and future headsets and smart glasses. We surmise that standardization in XR research will not arrive through specification alone. The empirical characterization reported above shows that even when a runtime behaves as its specification permits, the meaning of its data must be established empirically, and similar work will be needed across SDK versions, hardware revisions, and sensor configurations as they evolve. fMRIPrep (Esteban et al., 2019) is a useful design inspiration here, not a 1:1 model: a community-maintained preprocessing tool for fMRI. ResXR is released under Apache 2.0, keeping academic and downstream reuse open while preserving a contribution path.

## Supporting information

Supplementary Data 1 - ResXR column registry

## Acknowledgements

We thank Tal Zipori for valuable contributions for early versions of the research template.

## Declarations

### Ethics approval

This study was approved by the Ethics Committee of Tel Aviv University (reference number 0009858-3).

### Consent to participate

Informed consent was obtained from all individual participants included in the study.

### Consent for publication

Participants consented to the publication of their data.

### Data availability

The three demonstration sessions, the pipeline configuration file and the resulting raw and derivative BIDS datasets and quality reports (Bergstein, Barel, Shai Basson, et al., 2026d) are available in the ResXR Python pipeline repository (https://github.com/ResXR/resxr-python-pipeline, release v0.2.2, archived at https://doi.org/10.5281/zenodo.23562103). The recordings of the logging benchmark (Bergstein, Barel, & Schonberg, 2026) are available at Zenodo, https://doi.org/10.5281/zenodo.23678518 (see Code availability). The stress-test recordings and their ground-truth annotations (Bergstein, Barel, Shai Basson, et al., 2026b), on which the Evaluation rests, are available at Zenodo, https://doi.org/10.5281/zenodo.23668483.

### Code availability

The ResXR code is openly available under the ResXR GitHub organization (https://github.com/ResXR). The ResXR Unity Research Template (v0.2.1) is available at https://github.com/ResXR/resxr-unity-research-template and the Python analysis pipeline (v0.2.2) at https://github.com/ResXR/resxr-python-pipeline; both are released under the Apache License 2.0, and the pipeline version described here is archived on Zenodo (https://doi.org/10.5281/zenodo.23562103). The code that reproduces the stress-test results is deposited with the stress-test recordings (https://doi.org/10.5281/zenodo.23668483). The recordings of the logging benchmark and the code that computes every reported benchmark result are available at Zenodo, https://doi.org/10.5281/zenodo.23678518.

### Use of AI tools

The authors used Claude (Anthropic) to assist with drafting and editing parts of the manuscript text, writing and reviewing code, and checking references and reported values against the data. The authors reviewed, verified and approved all AI-assisted output and take full responsibility for the content.

### Open Practices Statement

The code of the ResXR Unity template, including the three demonstration paradigms, and of the ResXR Python pipeline is available at https://github.com/ResXR (see Code availability). The three demonstration sessions and the Motion-BIDS datasets and quality reports that the pipeline produced from them are available at https://github.com/ResXR/resxr-python-pipeline, and the stress-test protocols at https://github.com/ResXR/xr-validation-protocols (release v1.1.0, https://doi.org/10.5281/zenodo.23679224). The stress-test recordings, their annotations and the code that reproduces the stress-test results are available at Zenodo, https://doi.org/10.5281/zenodo.23668483, and the logging benchmark’s recordings and analysis code at Zenodo, https://doi.org/10.5281/zenodo.23678518. None of the reported studies were preregistered.

