## Supplementary Material for "ResXR: infrastructure for quality-controlled, reproducible studies of human behavior in Extended Reality"

### Contents

Supplementary Note 1. Custom tables: column annotations and their checks

Supplementary Note 2. API profiling, stress-test procedure and channel curation

Supplementary Note 3. Tethered PCVR sessions

Supplementary Note 4. Pipeline stages, input requirements and adding a tracking system or check

Supplementary Note 5. Unity template: Base Scene components, gaze rays and plain-text output

Supplementary Note 6. Logging benchmark

Supplementary Table 1. Tools in Table 1 and the code versions read

Supplementary Table 2. Column prefix-to-stream routing in the pipeline

Supplementary Table 3. Per-session quality-check summary

Supplementary Table 4. Sensor stress-test protocols and results

Supplementary Table 5. Quality signals and the columns they govern

Supplementary Data 1. ResXR column registry (separate CSV file)

### Supplementary Note 1. Custom tables: column annotations and their checks

Each annotation must supply a human-readable description and should supply a format specifying the data type (e.g., “number”, “integer”, “string”); where applicable, it also specifies units, a levels mapping of value to label for categorical columns, and minimum and maximum bounds. The annotations of all tables are written to one schema file per recording (<recording_id>_CustomTables.json). Two checks report annotation problems as errors in the log: in the Unity editor, after every compilation, a check reports any field without a [ColumnInfo] annotation or with a malformed levels entry, and any column name that two custom tables annotate differently, and at the end of each session a runtime check repeats these checks on the headset, also reports an empty description or an unrecognized format, and writes each error to the session’s ResXRDebugLogs file. By design, neither check prevents the table from being recorded; neither reports a missing format, although the pipeline discards the whole schema when any column lacks one (Supplementary Note 4).

### Supplementary Note 2. API profiling, stress-test procedure and channel curation

To identify the available data channels, we examined the OVRPlugin.cs source code and cataloged the accessible fields of the Node API and the Hand, Eye, Face and Body tracking APIs, with their status fields, validity flags and confidence scores, producing a registry of 1,460 candidate data columns (Supplementary Data 1).

*Stress-test procedure.* Before assessment, categorical labels (e.g., confidence levels reported as “Low” or “High”) were mapped to binary integers, and composite status strings (e.g., the HandStatus bitfield) were decomposed into one binary column per flag; the stress-test recordings store this composite string, whereas the current template writes the flags as separate columns at recording time (see Unity experiment template in the main text).

For the ground-truth event timeline, a researcher performing each choreographed protocol (Supplementary Table 4) noted the timestamps of intentional tracking manipulations (e.g., hand leaving the field of view, finger occlusion onset, headset removal). This produced a semantic time-map distinguishing intervals of valid user behavior from intervals of intentionally induced tracking failures.

### Supplementary Note 3. Tethered PCVR sessions

Three brief PCVR sessions (107, 113 and 147 s; 17 May 2026) were recorded by compiling the template as a Windows executable (Unity 6000.0.68f1, OVRPlugin 1.110.0) and connecting the headset to a desktop PC (Intel Core i7-13700; 128 GB DDR5 RAM; NVIDIA GeForce RTX 4080) via a tethered link. The sessions produced the continuous and face files with head, hand, eye and face tracking data; the body-tracking columns, enabled in these sessions, stayed empty.

### Supplementary Note 4. Pipeline stages, input requirements and adding a tracking system or check

This note describes the Python pipeline at version 0.2.2 and, where stated, the Unity template at version 0.2.1.

#### Requirements for data from other sources

These requirements were read from the pipeline code and tested on modified copies of a demonstration session (October 2026).

*Files.* When several files in the session folder match a pattern, the most recently modified one is used, with a warning. The custom-tables folder must be named {session_id}_CustomTables; each table's name is its CSV filename stem without the leading {session_id}_ prefix. CSV files must use commas as delimiters, a dot as the decimal separator and a header row; UTF-8 with or without a byte-order mark, either line ending and any column order are accepted.

*Clocks.* The timeSinceStartup column must be numeric, and its name is fixed in the code. Its first finite non-zero value is the session's time zero, and the face file's timeSinceStartup is assumed to be on the same clock. Clock values are never converted, and the rows of the continuous-data and face files are never sorted or interpolated. Each stream's own time axis is the column named for it under alternate_time_columns in the configuration; leading and trailing zeros in that column are allowed.

*Columns.* Channel type and units follow the name suffix, case sensitively: _px, _py, _pz, _x, _y, _z and HitPosition_X/Y/Z are positions in meters; _qx, _qy, _qz and _qw are quaternion components; _Time columns are latencies in seconds; validity suffixes such as _Present, _Valid_Position and _IsValid are boolean; columns ending in _Confidence, Confidence_Lower or Confidence_Upper and facial expression weights are typed MISC with units "normalized"; other columns, the hand and finger confidences among them, are typed MISC without units. Values are not converted. A stream is created only if its system is enabled in the configuration, its *_enabled key in the metadata file is true (Head excepted), and at least one column routes to it.

*Metadata.* The pipeline reads session_id (custom-tables folder and report name), hands_enabled, eyes_enabled, face_enabled, body_enabled and controllers_enabled, fixedDeltaTime, utc_start_iso8601 (acq_time in scans.tsv) and any key whose name contains "version" (SoftwareVersions in motion.json).

Departures that skip a session stop it with an error that names the cause, for example a missing or misnamed continuous-data or metadata file, a clock column without usable values, or an events file without an onset column; the other sessions continue. Configuration errors stop the run before any session is processed. The following departures pass with a logged error or warning that names them:

| Departure | What happens |
|---|---|
| timeSinceStartup or a stream's own clock that steps backward | The rows are kept in file order, so latencies decrease there and, for timeSinceStartup, flags next to those rows may be wrong. |
| Clock values in milliseconds (timeSinceStartup with a median step above 0.5 s, or a stream's own clock) | The values are read as seconds and not converted, so the latencies, sampling rates, flag times and event onsets that depend on them are wrong. |
| Empty or zero timeSinceStartup values | Their latency is n/a, and no quality flag or masking covers these rows. |
| A stream without its own clock | It uses timeSinceStartup as its time axis; its hand-tracking-loss, eye-closure, sampling-rate and clock-dropout checks raise no flags and are listed as not run. |
| A stream whose *_enabled key is missing or not true or false, a stream enabled without columns, or a missing or empty face file while face_enabled is true | The stream is not exported; without the face file, the eye-closure check is listed as not run. |
| Columns that no stream takes, including face columns placed in the continuous-data file | They remain only in the verbatim sourcedata copy. |
| Check input columns (HandTracked, Eyes_Closed) that are missing or renamed, or HandTracked values other than 0/1, true/false or blank | The check raises no flags and is listed as not run. |
| A misnamed events file or custom-tables folder | It is not read; events.tsv is built from the other source, or is not written if there is none. |
| A missing or invalid custom-tables schema file, including one in which a column entry lacks a description or a format | The schema is read all-or-nothing, so events.tsv is written but its custom columns are not described in events.json. |

The command-line exit code is 1 when the configuration has an error, a session was skipped or no session was found, and 0 otherwise, including runs in which a check did not run. A check that raises an error produces no flags, so with masking enabled it masks nothing, and the HTML report lists it.

**Quality-check defaults and flag records**

By default, the hand-tracking-loss check flags every interval in which a hand's HandTracked column (configurable under validation.settings.tracking_flags) is 0 or empty (it has no minimum duration); the eye-closure check flags intervals in which both Eyes_Closed_L and Eyes_Closed_R reach or exceed 0.9 for at least 0.1 s; the sampling-rate check flags an effective rate more than 10% from the expected rate, and irregular intervals when the coefficient of variation of the intervals between the stream's own samples exceeds 0.5; and the clock-dropout check flags zero or missing per-system timestamps between the first and last non-zero values. Each flag records its check, tracking system, extent on the application clock, severity, whether it may contribute to masking and optional target columns, without which it applies to the entire stream (columns governed: Supplementary Table 5).

**BIDS export**

For each tracking system, channels.tsv gives every column's BIDS channel type (LATENCY, POS, ORNT, or MISC), component, tracked point, units, sampling frequency and reference frame; channels.json the reference frames' rotation rule, rotation order and spatial-axis layout; and motion.json device and task metadata, channel counts by type, the nominal sampling frequency (the configured expected rate) and the effective one (note to Supplementary Table 3). Neither frequency is the rate of the rows: the template writes one row per fixed time step (0.01 s in the demonstration sessions), consecutive rows can carry the same per-system timestamp, and the time of each row is given by its latency channels. Each stream's per-system timestamp becomes a latency channel, in seconds from its own first finite non-zero value, and timeSinceStartup becomes latency_global, in seconds from the session's time zero, also the zero of events.tsv onsets; rows with a zero or non-finite clock value get NaN latency.

**Quality report**

(a) Session header and summary: manufacturer and model from the configuration file; platform, engine and SDK versions, sampling mode, fixed timestep, schema revision and the recorded systems, from the session record; total duration and warning count. (b) Per-stream statistics: row and channel counts, expected versus effective sampling rate, NaN % and warnings, expandable to per-channel descriptives. (c) Quality-flag table: check, stream, channel group, severity, onset, offset, duration and message of every flag. (d) Flag timeline, with instantaneous and duration events on a separate track.

**Adding a tracking system or check**

*Tracking systems.* A column with an existing system's prefix joins that system without any change. A new tracking system is added in code, where the set of systems (TrackingSystem in core/constants.py) and their column prefixes (SYSTEM_COLUMN_PREFIXES) are defined; the configuration needs its expected sampling rate and, if it has its own clock, an entry under alternate_time_columns, without which the sampling-rate and clock-dropout checks raise no flags. It counts as enabled whatever the session metadata contain and is then split, checked by the checks that run on every stream and exported without further changes. Data in a new separate file, like the face data, need further code in both components.

*Quality checks.* A check is a class with a name, a description, the streams it requires, the settings it reads under validation.settings, and a call that returns a list of quality flags (the ValidationCheck interface in validation/registry.py). An instance registered with register_check runs when its name is listed under validation.enabled_checks in the configuration: on every stream if it requires none, otherwise once per session when all its streams are present. It can flag any stream; its flags appear in the report and, when marked for masking, contribute to masking. A check defined outside the package runs only when the pipeline is called from a Python script that imports it first.

### Supplementary Note 5. Unity template: Base Scene components, gaze rays and plain-text output

*Base Scene components.* ResXRPlayer manages the participant's tracking (head, hands and finger joints), eye tracking (ResXREyeTracker), and controller and pinch input;

ResXRDataManager handles continuous data recording and export; and ResXRSceneManager governs scene transitions.

*Gaze rays.* Each frame, up to three raycasts are computed: the combined ray and, when enabled through the ResXRDataManager recording configuration (on in the default configuration), one per eye.

*Plain-text output.* Because Motion-BIDS specifies tab-separated text for motion data, channel descriptors and event timelines (Jeung et al., 2024), acquiring in text means the acquisition format and the archival format are the same kind of artifact, and the conversion between them splits and relabels columns by name rather than decoding a binary encoding.

**Supplementary Note 6. Logging benchmark**

The four runs were recorded on one Meta Quest Pro (Horizon OS 2.6.0, OVRPlugin 1.110.0, Unity 6000.0.68f1) on 10 September 2026. Two runs logged the 607-column default schema at 100 Hz (head and controller nodes, eyes, hands, gaze, system status and face; Body off). The application-side measurements came from a benchmark harness added to the template, later committed as 5952e48 on its data-performance-test branch and archived with the benchmark data (https://doi.org/10.5281/zenodo.23678518). Each run was analyzed over its task window, from the first task-start event to the last data row (37 to 61 s).

Mean frame time was 13.9 to 14.0 ms in all four runs. The 95th-percentile frame time was computed per second and averaged over the task window. The OVR Metrics Tool export's stale_frames_consecutive column, which Meta's documentation does not define, was 2 or more in 10 of the 60 seconds with logging and in 4 of the 51 seconds without. The logger used 159.7 ms of main-thread time per second: converting values to text accounted for 67.3%, the collectors, which query the runtime and fill each row, for 22.9%, and writing rows to the operating system for 5.2%. A second run of the same configuration measured 137 ms per second, so the logger's main-thread time varied by about 15% between runs, whereas the two runs' mean and 95th-percentile frame times differed by less than 0.1 ms. Device power was 1.1% higher with logging (9,590 vs. 9,486 mW).

Each run was one complete Maze Navigation session of at most 110 s, with one run per condition for the headset telemetry. Repeated frames rose during the logging run but not during the baseline run. In both pairs the logging run was recorded first.

**Supplementary Table 1**

*Tools in Table 1 and the code versions read*

| Tool | Version read | Commit | Code location | Read (2026) |
|---|---|---|---|---|
| ResXR Unity template (this paper) | v0.2.1 | – | https://github.com/ResXR/resxr-unity-research-template | 28 Oct |
| ResXR Python pipeline (this paper) | v0.2.2 | 2fa9aee | https://github.com/ResXR/resxr-python-pipeline | 27 Oct |
| ARETT (Kapp et al., 2021) | v2.0.0 (Unity toolkit) v1.0.0 (ARETT R package, gap fill) | d5927f7 cb1c459 | https://github.com/AR-Eye-Tracking-Toolkit/ARETT https://github.com/AR-Eye-Tracking-Toolkit/ARETT-R-Package | 4 Oct |
| LayerAwareLogging (Daralnakhla et al., 2026) | No release | 6feda80 | https://github.com/liris-xr/LayerAwareLogging | 2–5 Oct |
| EDIA (Klotzsche et al., 2025) | edia_core v0.6.1; edia_eye v0.6.0; edia_uxf v0.6.0; edia_lsl v0.6.1; edia_eye_quest no release (main branch) | edia_core 1fe2944 edia_eye 72ab48c edia_uxf 788a150 edia_lsl 8e73f1c edia_eye_quest bf64198 | https://github.com/edia-toolbox (five repositories) | 4–5 Oct |
| Cognitive3D SDK (Cognitive3D, 2026) | 2.5.0 | 1eb9384 | https://github.com/CognitiveVR/cvr-sdk-unity | 2–4 Oct |
| StudyFramework (Ehret et al., 2024) | branch 5.5; no tag | e7a1a00 | https://git-ce.rwth-aachen.de/vr-vis/VR-Group/unreal-development/plugins/unreal-study-framework | 3–4 Oct |
| VRrec (Kobylinski et al., 2026) | main branch | f58e69b | https://github.com/OPI-PIB/VRrec | 4 Oct |
| CLOVR (Segarra Martinez et al., 2024) | No release | b6905ac | https://github.com/xrtlab/clovr | 3–4 Oct |
| PLUME (Javerliat et al., 2024) | PLUME-Recorder v4.0.0-beta | 044ce25 | https://github.com/liris-xr/PLUME-Recorder | 3–4 Oct |
| UXF (Brookes et al., 2020) | 3.0.0 | 012103b | https://github.com/immersivecognition/unity-experiment-framework | 4 Oct |
| vexptoolbox (Schuetz et al., 2023) | v0.1.2 | 4d0c90d | https://github.com/ischtz/vizard-experiment-toolbox | 4 Oct |
| Ouvrai (Cesanek et al., 2024) | No release | 5b5ca35 | https://github.com/EvanCesanek/Ouvrai | 4–6 Oct |
| BeMoBIL pipeline (Klug et al., 2022) | v2.0.1 | 4161fd5 | https://github.com/BeMoBIL/bemobil-pipeline | 2–4 Oct |
| pymovements (Krakowczyk et al., 2023) | v0.28.0 | fe333f3 | https://github.com/pymovements/pymovements | 2–4 Oct |
| Motion-BIDS (Jeung et al., 2024) | BIDS specification v1.11.2 | fbe1af9 | https://github.com/bids-standard/bids-specification | 4 Oct |

*Note.* VRrec: the tagged releases (v1.0.0, v1.0.1) are on a separate history with different code; the R validation script that Table 1 lists is on the main branch only. PLUME: PLUME-Protos v3.0.0 (2eda315), PLUME-Viewer (cd27ed5) and PLUME-Python v1.3.2 (7577e1a) were also read.

### Supplementary Table 2

*Column prefix-to-stream routing in the pipeline*

| Column prefix or name | Signal |
|---|---|
| *Head* | |
| Node_Head_ | Head node (Node_Head_Time, the stream's time axis) |
| FocusedObject | Name of the object hit by the combined (cyclopean) gaze ray |
| RecenterCount, recenterEvent, shouldRecenter | Cumulative recenter count (OVRPlugin.GetLocalTrackingSpaceRecenterCount); recenter event (1 on the row where shouldRecenter changes from 0 to 1); recenter flag (OVRPlugin.shouldRecenter) |
| TrackingLost | Positional tracking lost (eye-center node not position-tracked) |
| UserPresent | Whether the user is wearing the headset, as reported by the SDK (OVRPlugin.userPresent) |
| TrackingOriginChange_ | Tracking-origin change event (OVRManager.TrackingOriginChangePending) and the pose of the new tracking origin relative to the previous one, passed through the template's world-space conversion |
| TrackingTransform_ | Pose returned by OVRPlugin.GetTrackingTransformRawPose, converted to world space by the template |
| timeSinceStartup | Application clock (Unity Time.realtimeSinceStartup); written as latency_global in every stream |
| *Hands* | |
| Node_HandLeft_, Node_HandRight_ | Hand nodes (Node_HandLeft_Time is the stream's time axis) |
| LeftHand_, RightHand_ | Hand status flags (HandTracked, InputStateValid, SystemGestureInProgress, DominantHand, MenuPressed), root pose, hand scale, hand and finger confidence, the RequestedTimeStamp and SampleTimeStamp fields; OVRPlugin.GetHandState |
| Left_XRHand_, Right_XRHand_ | Hand bone poses (26 bones per hand in the demonstration sessions) |
| *Eyes* | |
| LeftEye_, RightEye_, Eyes_Time | Per-eye gaze orientation (quaternion), validity and confidence (Eye Tracking API); Eyes_Time: eye-tracking SDK timestamp (the stream's time axis) |
| Node_EyeCenter_ | Eye-center node (orientation identical to the head node; Supplementary Table 4, Panel B) |
| EyeGazeHitPosition_ | World-space hit point of the combined gaze ray |
| LeftEyeGazeHitPosition_, RightEyeGazeHitPosition_, LeftFocusedObject, RightFocusedObject, HasLeftEyeHit, HasRightEyeHit | World-space hit point of each eye's gaze ray, name of the object it hit, and whether it hit an object[a] |
| *Face (separate face-expression file)* | |
| Face_ | Face_Status (the face state's validity flag) and Face_Time (SDK timestamp, the stream's time axis) |
| Brow_, Cheek_, Chin_, Dimpler, Eyes_Closed, Eyes_Look, Inner_Brow, Jaw_, Lid_, Lip_, Lips_, Lower_Lip, Mouth_, Nose_, Outer_Brow, Upper_Lid, Upper_Lip, Tongue_ | Facial expression weights (70 per sample) |
| FaceRegionConfidence | Upper- and lower-face confidence |
| *Optional streams (off in the shipped pipeline configuration)* | |
| Body_ | Body-tracking SDK time (Body_Time), confidence, fidelity, calibration status, skeleton change count, and joint poses with per-joint validity and tracked flags |
| Node_ControllerLeft_, Node_ControllerRight_ | Controller nodes |
| *No stream* | |
| Custom_ | Custom transforms recorded from the template inspector; kept only in the sourcedata copy |

*Note.* Columns are assigned to a stream when their name begins with one of the stream's prefixes (core/constants.py in pipeline v0.2.2); signals are those of the Unity template at version 0.2.1. Node_ columns hold the node's pose, presence, validity and tracked bits, and SDK time. The Face stream holds every column of the separate face-expression file, whatever its name; face columns placed in the continuous-data file are not exported. Face prefixes only mark facial expression weights as normalized MISC channels. Every column of the default schema begins with one of these prefixes; columns with no listed prefix, such as Custom_, are not exported.

[a]Recorded only when the template's per-eye gaze option (includeSeparateEyesGaze) is enabled, as it is in the default configuration.

**Supplementary Table 3**

*Per-session quality-check summary*

| Measure | sub-01 Binary Choice | sub-02 Maze Navigation | sub-03 Museum Viewing |
|---|---|---|---|
| Quality flags, all checks | 14 | 40 | 174 |
| Hand tracking loss | 9 | 28 | 67 |
| Sampling rate: rate deviation | 3 | 2 | 3 |
| Sampling rate: irregular intervals | 0 | 4 | 0 |
| Eye closure (Face and Eyes streams) | 2 | 6 | 104 |
| Clock dropout | 0 | 0 | 0 |
| Eye-closure flag episodes | 1 | 3 | 52 |
| Head effective rate, Hz (expected 72) | 71.0 | 66.9 | 68.1 |
| Hands effective rate, Hz (expected 90) | 71.6 | 67.2 | 68.2 |
| Eyes effective rate, Hz (expected 30) | 66.0 | 57.3 | 63.3 |
| Face effective rate, Hz (expected 30) | 35.6 | 31.2 | 35.3 |

*Note.* Values from the HTML quality reports of the example output committed with pipeline v0.2.2 (shipped configuration, masking off). In each session, the first hand-tracking-loss flag of each hand starts at the recording onset, because the hands are not tracked until the hand tracker's first sample. The effective rate is the number of distinct per-system timestamps from the first non-zero value, minus one, divided by the time they span.

### Supplementary Table 4

*Sensor stress-test protocols and results*

*Panel A. Protocols in the community registry, with results in the annotated recordings*

| Protocol | Result |
|---|---|
| Baseline (HAND-BASELINE) | As expected, HandTracked stayed 1 and hand confidence stayed High; InputStateValid dropped briefly in TEST_006. |
| Occlusion (HAND-OCCLUSION) | As expected, HandTracked dropped to 0 in every annotated event; hand position read 0, not the expected frozen last value. |
| Fist (HAND-FIST) | HandTracked stayed 1 (TEST_002, TEST_004) or dropped briefly (TEST_007); unlike the expected finger-tracking loss, per-finger confidence stayed High on all or nearly all tracked rows. |
| Finger Single Occlusion (HAND-FINGER-SINGLE-OCCLUSION) | Unlike the expected per-finger loss, the covered left hand's five finger-confidence columns stayed High on all or nearly all tracked rows; HandTracked stayed 1 except for a few seconds in both hands shortly before TEST_007 ended. |
| Tracking Boundary (HAND-TRACKING-BOUNDARY) | HandTracked dropped to 0 in every annotated boundary event; unlike the expected sequence, hand confidence went Low when HandTracked dropped, not before. Node API Valid_Position and both Tracked bits read 0 during the losses; Valid_Orientation stayed 1. |
| Occlusion (EYE-OCCLUSION) | Unlike the expected drop, LeftEye_IsValid and RightEye_IsValid stayed 1. |
| Both eyes closed (EYE-BOTH-EYES-CLOSED) | Unlike the expected drop, LeftEye_IsValid and RightEye_IsValid stayed 1; the eye-closed weights detected the closures (TEST_003, TEST_008). |
| Gaze Boundary (EYE-GAZE-BOUNDARY) | Unlike the expected drop or degradation, LeftEye_IsValid and RightEye_IsValid stayed 1 at the gaze extremes, and eye confidence held one fixed value per recording whenever IsValid was 1. The eye-closure check flagged the eyes as closed at the downward extreme (TEST_003, TEST_008, TEST_009). |
| Gaze Boundary Closed (EYE-GAZE-BOUNDARY-CLOSED) | Unlike the expected drop, LeftEye_IsValid and RightEye_IsValid stayed 1; as expected, the eye-closed weights detected the closures (TEST_003, TEST_008). |
| Neutral baseline (FACE-NEUTRAL-BASELINE) | As expected, nearly all expression weights stayed below 0.1 and both face-region confidences stayed near 1. |
| Blink (FACE-BLINK) | Both eye-closed weights rose to 0.9 or more with the blinks: 40 brief rises, each 0.2 s or shorter, in about 10 s of rapid blinking (TEST_003). |
| Lower Face Occlusion (FACE-LOWER-FACE-OCCLUSION) | As expected, lower-face confidence fell and upper-face confidence stayed 1; Face_Status stayed true. |
| Headset Removal (HEAD-HEADSET-REMOVAL) | Unlike the expected drop, the head node's Valid and Tracked flags stayed 1 while the headset was off, as did LeftEye_IsValid and RightEye_IsValid; Face_Status stayed true. UserPresent, which the current template records by default (Supplementary Table 2), was not recorded in these sessions. |
| Arm Swing (HAND-ARM-SWING) | Repeated short HandTracked losses in every annotated window, where none were expected; the longest 1.2 s. |
| Head Rotation, Fixed Gaze (EYE-HEAD-ROTATION-FIXED-GAZE) | As expected, LeftEye_IsValid and RightEye_IsValid stayed 1. |
| Sustained Expressions (FACE-SUSTAINED-EXPRESSIONS) | Face_Status stayed true, as expected; for each instructed movement, at least one expression weight named for it rose from near 0; unlike the expected high confidence, lower-face confidence fell while the tongue was held out (TEST_009). |
| One eye closed (EYE-ONE-EYE-CLOSED) | Unlike the expected drop, LeftEye_IsValid and RightEye_IsValid stayed 1; the closed eye's eye-closed weight rose to a median of 0.63 to 0.74, short of the expected value near 1, and the open eye's stayed near 0 (TEST_003, TEST_008). |

*Panel B. Cross-API checks on the stress-test recordings*

| Check | Recordings | Result |
|---|---|---|
| Node API hand pose vs. Hand API root pose | All 11 | Identical, to the 7 significant digits the template writes, on every hand-tracked row; on lost rows both positions read 0. |
| Hand-node velocity | All 11 | Velocity and angular velocity were 0 on every row, including while the hand was tracked and moving; head- and eye-node velocities were populated. |
| Body API during induced hand loss | TEST_004, TEST_006, TEST_010 | The lost side's hand and arm joints never read zero: they moved to the same fixed arm pose while flagged as valid, and only their Tracked flags marked the loss. Body confidence, fidelity, calibration status and skeleton-change count stayed constant. |
| Body wrist vs. Hand API wrist while tracked | TEST_004 to TEST_011b | Agreed closely but were almost never identical. |
| Node eye orientation vs. head orientation | All 11 | Equal on every row (per-eye nodes to within rounding); during eye-only gaze shifts, the Eye API's gaze moved and the node eye moved only with the head. |

*Note.* The community registry (https://github.com/ResXR/xr-validation-protocols, release v1.1.0, https://doi.org/10.5281/zenodo.23679224) gives each protocol in full, including its expected behavior; IDs omit META-QUESTPRO-. All rows: Meta Quest Pro; SDK / Firmware "Meta XR SDK v78 / fw v79-v83". The recordings and their annotations are available at Zenodo, https://doi.org/10.5281/zenodo.23668483. An event counted as detected if a flagged interval (Supplementary Table 5) overlapped its annotated window within 2 s. Template builds 2.5 and 2.7 stored the two face-region confidence columns swapped. TEST_003's face data come from the earlier logger, recorded in the same session. Annotation times were noted in whole seconds, on a clock inferred from their time ranges.

**Supplementary Table 5**

*Quality signals and the columns they govern*

| Check | Stream | Signal and rule | Severity | Columns governed |
|---|---|---|---|---|
| Hand tracking loss (left and right) | Hands | Each hand's HandTracked column (LeftHand_Status_HandTracked, RightHand_Status_HandTracked) 0 or empty; no minimum duration | warning | The hand's position, orientation, scale and confidence columns (left: Node_HandLeft_ and LeftHand_Root_ pose, Left_XRHand_ bone poses, LeftHand_HandScale, LeftHand_HandConfidence and the five LeftHand_FingerConf_ columns; right: the same with Node_HandRight_, RightHand_ and Right_XRHand_), 203 per hand in the demonstration sessions. Presence, validity, tracked, status and timing columns are kept. |
| Eye closure (source) | Face | Eyes_Closed_L and Eyes_Closed_R both ≥ 0.9 for ≥ 0.1 s | info | None: the flag names Eyes_Closed_L and Eyes_Closed_R, the columns it reads |
| Eye closure (propagated) | Eyes | Eyes-stream rows within each Face-stream eye-closure interval | info | Per-eye gaze orientation (LeftEye_ and RightEye_ quaternions) and gaze-hit positions (EyeGazeHitPosition_, LeftEyeGazeHitPosition_, RightEyeGazeHitPosition_); 17 columns in the demonstration sessions. The Node_EyeCenter pose and the validity and confidence columns are kept. |
| Clock dropout | Each stream | Zero or missing per-system timestamp between the first and last non-zero values | warning | All data columns of the stream in the affected rows |
| Sampling rate | Each stream | Effective rate more than 10% from the expected rate; coefficient of variation of the intervals between the stream's own samples (its SDK timestamps) above 0.5 | warning | None (advisory) |

*Note.* Masking is off by default (apply_quality_masking: false in the shipped configuration). When it is enabled, every flag marked for masking contributes unless masking_checks lists a subset; masked values are written as n/a, and the latency and latency_global columns are never masked. Thresholds, and the flag columns the hand-tracking-loss check reads, are configurable. Source: pipeline v0.2.2 code and a run of the three demonstration sessions with masking enabled (27 October 2026).

### Supplementary Data 1. ResXR column registry

Supplementary_Data_1_ResXR_column_registry.csv (separate file) catalogs the 1,460 candidate data columns. Each row gives the column name, the file it belongs to (dataset: continuousdata or faceexpressions), its API source (api), comments, and whether it is in the default schema (exclusion_status: 607 included; 853 excluded, 777 of them the body-tracking columns that the template writes when its Body option is enabled).
